# Mechanism of Phosphate Release from Actin Filaments

**DOI:** 10.1101/2023.08.03.551904

**Authors:** Yihang Wang, Jiangbo Wu, Vilmos Zsolnay, Thomas D. Pollard, Gregory A. Voth

## Abstract

After ATP-actin monomers assemble filaments, the ATP’s γ-phosphate is hydrolyzed within seconds and dissociates over minutes. We used all-atom molecular dynamics simulations to sample the release of phosphate from filaments and study residues that gate release. Dissociation of phosphate from Mg^2+^ is rate limiting and associated with an energy barrier of 20 kcal/mol, consistent with experimental rates of phosphate release. Phosphate then diffuses in an internal cavity toward a gate formed by R177 suggested in prior computational studies and cryo-EM structures. The gate is closed when R177 hydrogen bonds with N111 and is open when R177 forms a salt bridge with D179. Most of the time interactions of R177 with other residues occludes the phosphate release pathway. Machine learning analysis reveals that the occluding interactions fluctuate rapidly, underscoring the secondary role of backdoor gate opening in P_i_ release, in contrast with the previous hypothesis that gate opening is the primary event.

**Significance Statement:** The protein actin assembles into filaments that participate in muscle contraction and cellular movements. An ATP bound to the actin monomer is hydrolyzed rapidly during filament assembly, but the γ-phosphate dissociates slowly from the filament. We identified phosphate dissociation from Mg^2+^ as the rate-limiting step in phosphate release from actin based on an energy barrier that aligns with the experimentally determined release rate. The release of phosphate from the protein requires opening a gate in the actin molecule formed by the interaction between sidechains of arginine 177 and asparagine 111. Surprisingly, simulations revealed other interactions of the sidechain of arginine 177 that occlude the release pathway most of the time but have not been observed in low-temperature cryo-EM structures.

## Introduction

Living cells recycle actin molecules countless times by assembly and disassembly of filaments that they use for cellular movements as well as physical support of the cell and its plasma membrane. During each cycle of assembly, actin molecules with bound Mg-ATP from the cytoplasm (1) incorporate at one of the two ends of a filament (largely the fast-growing “barbed” end) (2, 3) and undergo a conformational change that flattens the twisted actin monomer (4-6). Flattening repositions residues in the active site and activates the ATPase activity of the subunit about 42,000-fold (7, 8) with a half time of 2 s, producing ADP and an H_2_PO_4_^-^ ion, henceforth called phosphate (P_i_), to form an intermediate, Mg-ADP-P_i_-actin. The limiting cases for the reaction occur by either a dissociative pathway, where the ß-γ-phosphate bond breaks prior to the addition of lytic water, or a concerted mechanism, where the γ-phosphate bond breaks concurrently with the addition of lytic water (9-13). The most state-of-the-art computer simulations via quantum mechanics/molecular mechanics (QM/MM) with biased molecular dynamics free energy sampling point to a mechanism in between these two limits (11, 12). Although hydrolysis releases free energy, the structures of ATP- and Mg-ADP-P_i_-actin subunits are identical (6). The free γ-phosphate escapes into solution very slowly with a half time of 6 min (14-16). Filaments composed of ADP-actin subunits differ those with ATP or ADP-P_i_-actin subunits in stiffness (17-21), have higher affinities for regulatory proteins such as cofilin (22-25), and differ in the response to stresses (26, 27).

Despite ever higher resolution structures of actin filaments, now extending to < 2.5 Å resolution (28, 29), the mechanism of slow phosphate release is still in question. Wriggers and Schulten (30) concluded from steered molecular dynamics (SMD) simulations that phosphate is released from monomeric actin by a pathway from the active site to a “backdoor” near methylhistidine HIC73 in subdomain 1 (SD1). They limited their analysis to an assumed a predefined pathway and applied an external force to pull phosphate along to exit actin. Along this path the phosphate interacts with several residues including S14, G74, G158, H161 and V159. The rate limiting step in their simulations was dissociation of phosphate from the divalent cation in the active site, assumed to be Ca^2+^ rather than the physiological Mg^2+^. However, as pointed out by more recent studies (31-33), SMD can produce slow-converging, significantly out of equilibrium trajectories, potentially less representative of physiological molecular dynamics. Cryo-EM structures of actin filaments with bound AMPPNP (an ATP analog)-, ADP-P_i_-, and ADP-actin filaments nucleotides suggested a possible exit gate formed by R177, N111, and HIC73, which was closed in the ADP-P_i_-actin structure and open in the ADP-actin structure. A possible mechanism is that as phosphate dissociates from the Mg^2+^ ion, a hydrogen bond (H-bond) between S14 and G74 breaks making SD2 more flexible and favoring rotation of the side chain of HIC-73 to open the exit gate. On the other hand, the N111-R177 exit gate was closed in higher resolution cryo-EM structures of ADP-P_i_-, and ADP-actin filaments (28, 29).

We have employed well-tempered metadynamics (WT-MetaD) (34-37) biased free energy sampling and unbiased molecular dynamics (MD) simulations to investigate the release of P_i_ from Mg-ADP-P_i_-actin filaments. We find a free energy barrier of 20.5 kcal/mol associated with breaking the bond between Mg^2+^ and the H_2_PO_4_^-^ ion, which agrees with the experimental time scale for P_i_ release according to a simple transition state theory estimate. Once free from Mg^2+^, P_i_ diffuses through an internal cavity to a gate formed by the side chain of R177 in subdomain 3. Different hydrogen bonding interactions of R177 characterize the three main states of the channel connected to the exterior. Formation of a hydrogen-bond between the side chains of R177 and N111 in subdomain 1 closes the gate. Alternately, the sidechain of R177 can bend toward the interior of the protein and form a fluctuating network of hydrogen bonds with H73, H161 and P109, which also occludes the channel. Formation of a salt bridge with D179 opens the gate on the channel. During unbiased MD simulations, actin spends most of the time in the occluded state but also visits the closed and open states. Time in the occluded and closed states slows phosphate release, but it is not rate-limiting in our simulations.

## Results

### Preparation of a pentameric ADP-P_i_ actin filament for MD simulations

We used both WT-MetaD and unbiased MD simulations to investigate the release of P_i_ from a subunit in a short, ADP-P_i_-actin filament with five subunits named from the barbed end A0-A4 (Fig. 1A). We focused on the middle subunit A_2_ within the accessible time in all the simulations. We prepared these pentamers as follows:

**Figure 1.**
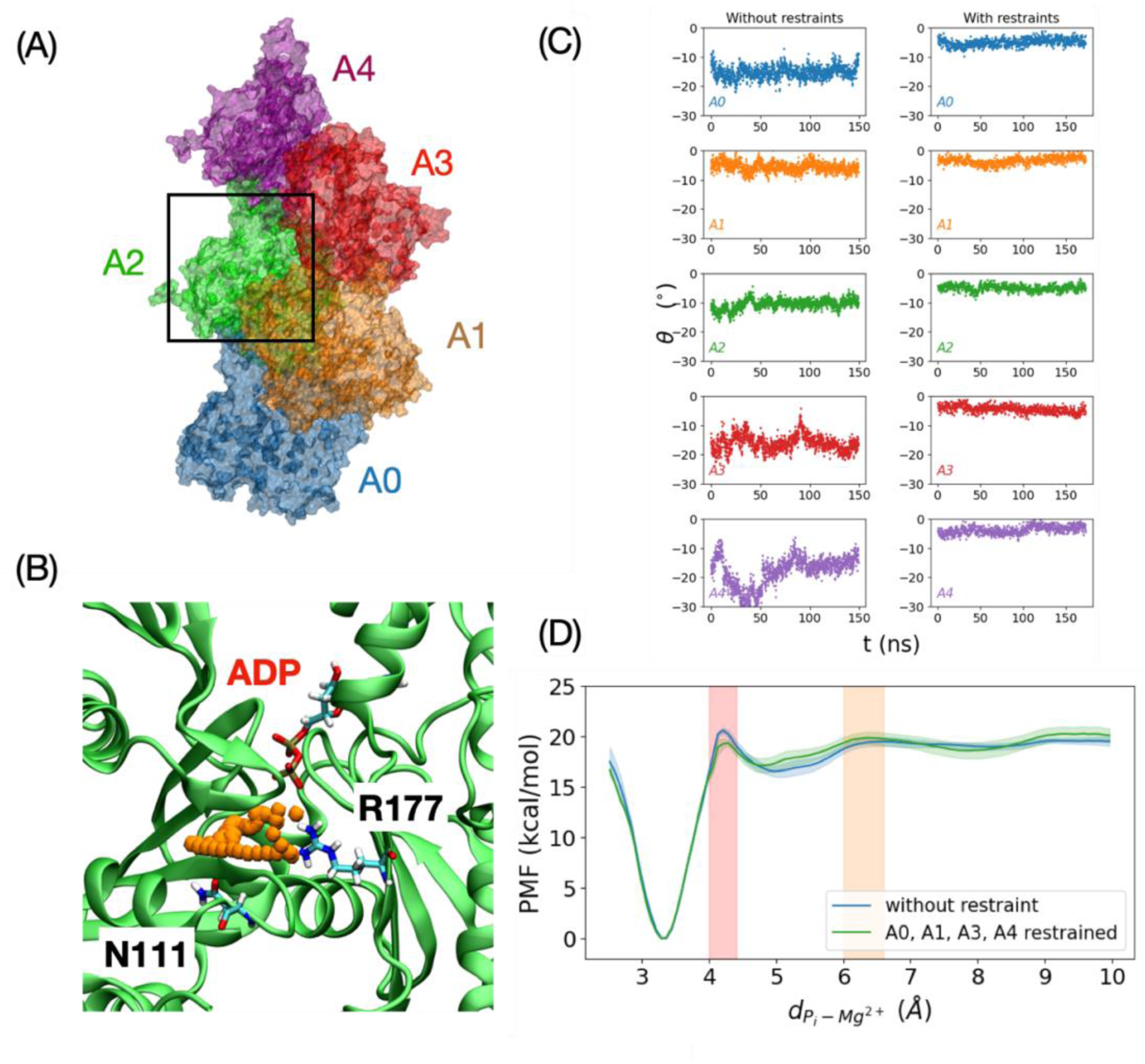
Pathways of P_i_ release in independent WT-MetaD simulations. (Panel A) Cartoon representation of the actin filament 5-mer with the subunits colored in blue, orange, green, red, and purple from A0 at the barbed end to A4 at the pointed end. Our simulations focus on the phosphate release from actin subunit A2, which is highlighted by the black box. (Panel B) One representative trajectory of phosphate release (orange spheres) from actin subunit A2 (green cartoon representation) made by plotting the positions of the phosphorus in each frame. R177 and N111 indicate the exit of the “backdoor” pathway (see Supplementary Fig S1 for details of other Pi release trajectories). (Panel C) Time course of the subunit dihedral angles in WT-MetaD simulations with unrestrained simulations in the left column and restrained simulations in the right column with a force constant of 5000 kJ/mol/radian^2^ applied on subunits A0, A1, A2 and A4. (Panel D) Potential of Mean Force (PMF) as a function of the distance between phosphate and the Mg^2+^ ion from the constrained (green) and unconstrained (blue) simulations. The shaded regions are standard errors. The positions of the energy barriers are highlighted by light orange and red.

Starting with a 13-subunit section of the Chou (6) cryo-EM structure of the ADP-P_i_-actin filament (PDB 6DJN), Zsolnay et al. carried out computational solvation and equilibration (38), which produced a filament with altered subunits on the ends and a core of conventional subunits in the occluded state (defined below) with helical parameters similar to the cryo-EM structure.

We took five conventional subunits from the core of the 13-mer filament and further equilibrated the system with energy minimization and a 10 ns simulation with a constant number of particles, pressure and temperature (NPT) ensemble at temperature 310 K and pressure 1 bar **(Fig 1A)**. This resulted in flattening of the terminal subunits at the barbed end and partially twisting of the terminal subunits at the pointed end. The central subunit A2 was flattened with an occluded phosphate release pathway, described in detail below. We used this structure as the initial structure in the unconstrained MD simulations.

The altered conformations of the subunits flanking subunit A2 may influence the release of phosphate, so we carried out four additional constrained simulations with restraints to maintain the flattened state of the terminal subunits flanking the central subunit. The initial structure for the constrained simulations was prepared by using steered MD to drive all the subunits into the flattened state (as detailed in the Methods section, with subsequent WT-MetaD results given in the **Supplementary Material**).

A recent 2.22 Å resolution structure of the Mg-ADP-P_i_-actin filament (PDB 8A2S) has water molecules resolved in the active site, allowing for comparison with our hydration of the lower resolution structure. Starting with five subunits of PDB structure 8A2S, we followed the same protocol to set up the system and perform the WT-MetaD simulations. The two hydrated structures were similar, so the following text focuses on simulations based on the PDB 6DJN with comparisons of simulations starting with the PDB 8A2S structure.

### Well Tempered-MetaD simulations

In six independent WT-MetaD simulations of unrestrained filaments and four simulations of restrained filaments, phosphate used similar pathways to move from the active site next to the ß-phosphate of ADP out of the protein next to R177, commonly called the "backdoor" pathway **(Fig. 1B**). Thus, the more stable dihedral angles of all five subunits in the restrained pentamers did not have a strong influence on phosphate release, despite the central subunit A2 being more flattened in the restrained pentamer **(Fig. 1C)**. Furthermore, the potentials of mean force (PMFs, i.e., free energy curves) as a function of the distance between P_i_ and the Mg^2+^ ion were similar in the restrained and unrestrained filaments **(Fig. 1D)**. Additionally, the restraints had little or no effect on the relative stability of the R177 backdoor gate conformations **(Supplementary Fig. S2)**. The rest of the results section therefore focuses on simulations with unrestrained subunits but also notes similar results of simulations with retrained subunits.

### The potential of mean force as phosphate is released by the “backdoor” pathway

We calculated the PMF as a function of the distance between P_i_ and the Mg^2+^ ion in the active site by averaging the free energy profiles of 6 independent unbinding trajectories for both the unrestrained and restrained filaments (**Fig. 1D)**. The error bars in **Fig. 1D** are standard errors representing the uncertainty in the PMF measurements. The small variability indicates consistency between different simulation trials, whose energy profiles are separately shown in **Supplementary Fig. S1**. The small variance between individual release trajectories and PMF also suggests that the P_i_ release pathway is stable. Moreover, the gentle biasing potential that we added to the system did not introduce any artifact or spurious mechanism in the simulations.

The plot in **Fig. 1D** reveals two energy barriers in the release pathway: a high barrier of ∼20.5 kcal/mol close to the bound state, and another of ∼5.0 kcal/mol near the unbound state. The difference between the PMFs from unrestrained and restrained systems is within the simulation error. Based on the positions of these energy barriers in the PMF, phosphate proceeds in three distinct stages.

### Stage 1, leaving the bound state

The deep energy minimum for phosphate bound to Mg^2+^ in the nucleotide-binding site indicates strong interactions between phosphate and Mg^2+^ that are reinforced by interactions with the protein. P_i_ hydrogen bonds with S14 and Q137 in the initial equilibrium structure (Fig. 2D) and occasionally forms additional H-bonds or salt bridges with nearby residues A108, D154, and H161 during unbiased MD simulations.

**Figure 2.**
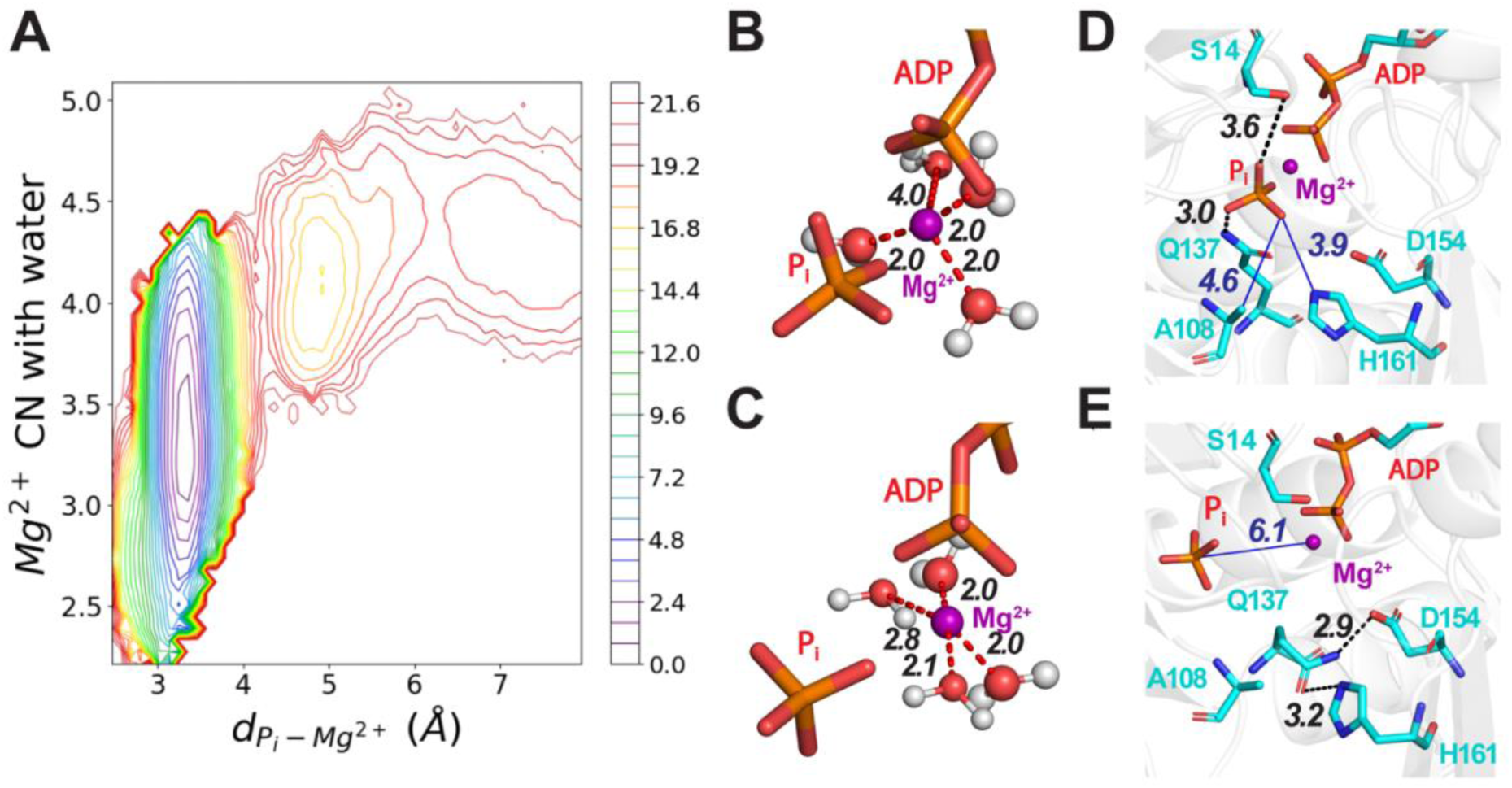
The release of P_i_ proceeds with the solvation of Mg^2+^ and the disruption of interactions with nearby residues near the active site. (Panel A) A 2-dimensional plot of potential of mean force (PMF) as a function of the number of waters coordinated with Mg^2+^ and the distance between P_i_ and Mg^2+^ calculated from one release trajectory. The bar on the right has the color code for energy levels in kcal/mole. Supplementary Materials Fig. S4 has PMFs from five other trajectories. B-E, Stick figures of the active site with Mg^2^ depicted as a purple sphere using the VDW presentation with a 0.8 scale, hydrogen bonds as black dashed lines, electrostatic interactions as red dashed lines, and nonbonding distances as blue solid lines. B-C, ADP and P_i_ at two time points during a WT-MetaD simulation. (Panel B) Mg^2+^ is coordinated with 3 water molecules. (Panel C) Mg^2+^ coordinated with 4 water molecules after losing coordination with P_i_. D-E, Hydrogen-bonds at two time points during a WT-MetaD simulation. (Panel D) Before dissociating from Mg^2+^, P_i_ was coordinated with the oxygen of the side chain of residue S14 and a hydrogen atom of the side chain amine group of residue Q137. (Panel E) After changing its rotameric conformation, the side chain of Q137 forms a hydrogen bond between its side chain oxygen and a hydrogen atom on the ring of H161. During our MD simulations, the reorientation of Q137 always occurs before P_i_ loses its coordination with Mg^2+^ (see Supplementary Materials for more details).

The release of phosphate begins with the loss of coordination between phosphate and the Mg^2+^ ion and disruption of interactions with nearby residues. These changes are unfavorable as reflected in the energy barrier at small distances from the bound state (**Fig. 1D**, pink shaded region). During WT-MetaD simulations, Q137 reoriented away from P_i_ with the distance between them becoming larger than 4.0 Å. After the disruption of its interaction with P_i_, Q137 transited into a new rotameric state (**Fig 2E**) to hydrogen bond with the side chain of H161. The distance between P_i_ and Mg^2+^ increased only after disruption of the interaction between Q137 and P_i_ and usually accompanies the forming of a hydrogen bond between Q137 and H161. (**Supplementary Fig. S2 and Supplementary Video 1)**. The conformational changes of other residues near the active site do not exhibit a direct correlation with the dissociation of P_i_ from Mg^2+^, typically retaining their configurations both before and after the event.

As phosphate moves away from Mg^2+^ in both unbiased and WT-MetaD simulations the number of waters coordinated with Mg^2+^ fluctuated in a range from 2.3 to ∼ 4.2 (**Figs. 2A and S3**) as estimated using a continuous switch function with a 3 Å cutoff (see **Supplementary Materials** for more details). **Figs. 2B** and **2C** illustrate times during the simulations where Mg^2+^ coordinated with three or four water molecules.

The PMF calculated from WT-MetaD trajectories revealed the relationships between the P_i_-Mg^2+^ distance and the number of waters coordinated with Mg^2+^ **(Fig. 2A**; energy scale 0.8 kcal/mol). The deep energy well in PMF profile at a P_i_-Mg^2+^ distance of 3.4 Å with 3.2 waters coordinated with Mg^2+^ indicates the most stable bound structure for P_i_. Above the high energy barrier at separations greater than 4.3 Å, phosphate and Mg^2+^ dissociate with 4-5 waters around Mg^2+^. This mechanism was observed in all simulated trajectories (**Supplementary Fig. S4**).

The locations of the phosphorus atom of the γ-phosphate during simulations of its release form three clusters in the cavity (**Figures 3A** and **3B)**. The most heavily populated cluster is close to Mg^2+^ and reflects the strong interactions that stabilize the bound state, aligning with the deep energy basin in the PMF (**Fig. 1 D)**. The other two clusters correspond to the two stages explained in following sections. The empty/sparse areas between the clusters correspond to the energy barrier regions (**Fig. 1 D)**.

**Figure 3.**
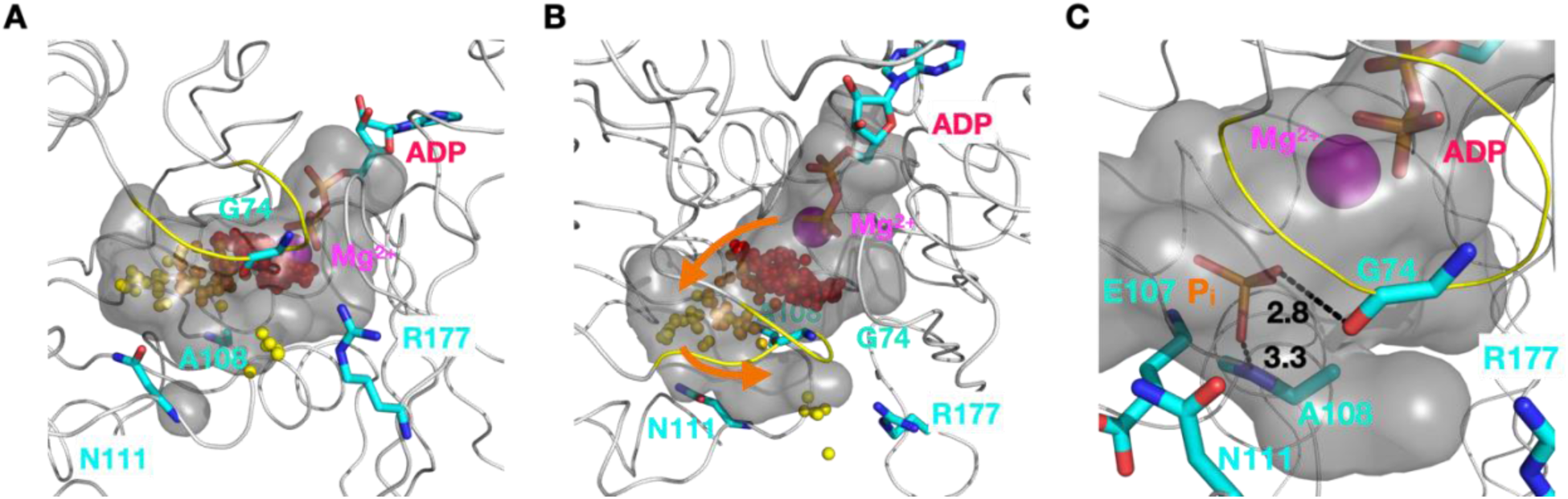
P_i_ diffuses in a cavity inside an actin subunit between the active site and the R177 back door in the occluded state. The cavity is shown as a grey surface defined by probing an equilibrium structure from an unbiased MD simulation in the occluded state with a detection radius of 4.2 Å. Mg^2+^ is shown as a purple sphere, ADP, G74, A108, N111, R177 are shown in the licorice representation. The S-loop (residues 70 to 79) is highlighted in yellow. The positions of the phosphorus atom of the γ-phosphate at 0.5 ns intervals during in a WT-MetaD simulation are shown as spheres with a gradient of colors ranging from red at early times to yellow at late times. (Panels A-B) Cavity from two viewpoints in the occluded state with orange arrows to indicate the path of the release process. Clusters are present next to Mg^2+^ and in the cavity leading up to the gate. (Panel C) During the release process, hydrogen bonds (black dash lines) form transiently between P_i_ (orange stick figure) and the backbone carbonyl oxygen of residue G74 and the amide hydrogen of residue A108.

### Stage 2, diffusion in the cavity

After dissociating from Mg^2+^, P_i_ diffuses in a cavity (**Fig. 3B**), which we delineated by probing with a detection radius of 4.2 Å, comparable in size to phosphate (**Figures 3A** and **3B)**. ADP and Mg^2+^ occupy one end of the “C”-shaped cavity with the R177 gate at the other end (**Fig. 3B**). In between phosphate diffuses between the sensor loop (residues 71–77) and residues E107 and A108, where it can form hydrogen bonds with the backbone atoms of G74 and A108 (**Fig. 3C)**. The locations of the phosphorus atom of phosphate in this diffusion process form the second largest cluster in Fig. 3A and 3B. The relatively flat free energy profile after P_i_ dissociates from Mg^2+^ indicates a lack of strong interactions during this stage (**Fig. 1C**).

### Stage 3, opening the backdoor

To exit the actin subunit, P_i_ must overcome a second barrier thought to be formed at least in part by the side chain of R177. In cryo-EM structures (PDB 6DJN, 8A2S, 8D17), the side chain of R177 forms hydrogen bonds with the side chains of N111 and H161, and the backbone of P109, creating a barrier to P_i_ release **(Fig. 4A**, **Table 2)**. This has been called the R177 closed state. Crystal structures of actin monomers (39) and a cryo-EM structure of the pointed end of the ADP-actin filament (40) have the gate open with a salt bridge between R177 and the sidechain of D179 but no hydrogen bonds with N111 or P109 **(Fig. 4E**, **Table 2)**. This is the open state.

**Figure 4.**
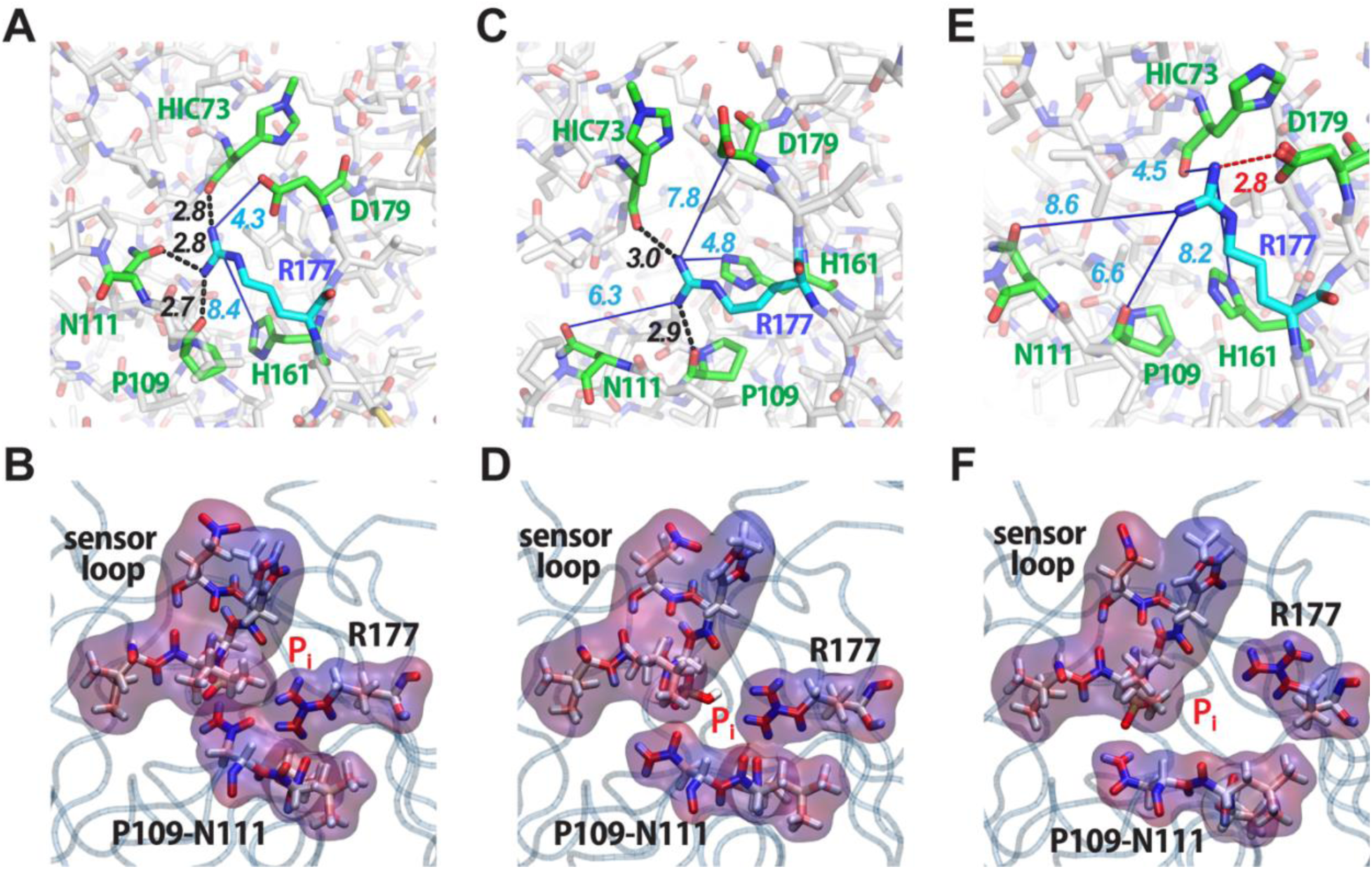
Hydrogen bonding networks of R177 in the (Panels A-B) closed state, (Panels C-D) occluded state and (Panels D-E) open state. The top row shows stick figures with hydrogen bonds as black dashed lines and longer distances to potential hydrogen bond partners as blue lines. The second row shows surface representations of the residues forming the gate in MD simulations. The surfaces of the sensor loop (residues 71 to 77), R177, N111, and P109 are colored by the charge density with red for positive charges and blue for negative charges. (Panel A) Cryo-EM structure of ADP-P_i_-actin filaments (PDB 8A2S) with a hydrogen bonds between R177 and the sidechain of N111 as well as the backbones of HIC73 and P109 closing the gate. (Panel B) Surface representation of the closed state in an MD simulation. (Panels C-D) Snapshots from an unbiased MD simulation in the occluded state. R177 hydrogen bonds with the backbone of P109 and the side chain of H161 precluding the release of phosphate. (Panels E-F) Open state with an electrostatic interaction between the sidechains of R177 and D179 that opens a channel for phosphate to escape. (Panel E) Crystal structure of MAL-RPEL2 complexed to G-actin (PDB 2V52) in the open state. (Panel F) Snapshot from a WT-MetaD simulation in the open state.

**Table 1.** Computer simulations performed in this study.

| Simulation | Biasing method | Initial state | Number of simulations | Special Settings |
| --- | --- | --- | --- | --- |
| P<sub>i</sub> release without constrained subunits | WT-MetaD | Occluded state | 6+4 | 6 set up with PDB 6DJN<br>4 set up with PDB 8A2S |
| P<sub>i</sub> release with constrained terminal subunits | WT-MetaD,<br>Harmonic restraint on the subunit dihedrals | Occluded state | 4 | - |
| Unbiased MD | - | Occluded state | 4+4 | 4 set up with PDB 6DJN<br>4 set up with PDB 8A2S |
| Low temperature 1 | - | Closed state | 1 | T=77 K |
| Low temperature 2 | - | Closed state | 1 | Lowering the temperature from 322 K to 77 K |
| Low temperature 3 | - | Closed state | 1 | Alternating the temperature between 322 K and 77 K |
| Low temperature 4 | - | Occluded state | 1 | Alternating the temperature between 322 K and 77 K |

**Table 2.** Distances between the sidechain of R177 and potential hydrogen-bonding partners in structures of actin monomers and filaments and in MD simulations. Distances smaller than 4.0 Å are highlighted in red. Measurements were taken from the following structures. Monomer: PDB 2V52. Internal: PDB 8A2T. P-1: Chou and Pollard. https://biorxiv.org/cgi/content/short/2023.05.12.540494v1. Barbed end: PDB 8F8R.

| Filament subunits (except monomer) | Nucleotide | R177-N111 SC | R177-P109 C=O | R177-H161 SC | R177-H73 C=O | R177-D179 SC | Gate |
| --- | --- | --- | --- | --- | --- | --- | --- |
| Monomer | ATP | 8.6 Å | 6.6 Å | 5.9 Å | 4.8, 4.5 Å | 3.2, 3.5 Å | Open |
| Internal | ADP | 2.9, 3.0 Å | 2.5 Å | 7.2 Å | 2.9 Å | 3.8 Å | Closed |
| Pointed end P-1 | ADP | 9.6, 10.6 Å | 6.8 Å | 5.5 Å | 3.3 Å | 3.0 Å | Open |
| MD closed | ADP-P<sub>i</sub> | 2.8 Å | 2.8 Å | 4.20 Å | 3.5 Å | 8.4 Å | Closed |
| MD occluded | ADP-P<sub>i</sub> | 8.2 Å | 4.0 Å | 5.1 Å | 4.9 Å | 7.4 Å | Occluded |
| MD open | ADP-P<sub>i</sub> | 10.3 Å | 6.5 Å | 7.3 Å | 4.9 Å | 2.79 Å | Open |
| MD wide-open | ADP-P<sub>i</sub> | 16.1 Å | 10.8 Å | 8.6 Å | 9.9 Å | 3.2 Å | Wide-open |

Our WT-MetaD and long unbiased MD simulations showed that the hydrogen-bonding interactions of R177 with HIC73, P109, N111, H161 and D179 fluctuate on a nanosecond (ns) time scale, visiting not only the open and closed states defined above but also a wide-open state and multiple configurations that also block phosphate release without a hydrogen bond between R177 and N111 **(Figs. 4C-D**, **Table 2)**. We call these blocking configurations occluded states. We used free energy profiles **(Fig. 5C)** to choose the cutoffs for the bonding distances to define four states of the phosphate gate based on distances between the side chains. The distance between two residues is determined as the distance between the closest pair of atoms involved in the formation of a hydrogen bond or a salt bridge. Table 2 lists the specific atoms participating in these interactions, and Supplementary Materials have detailed definitions.

**Figure 5.**
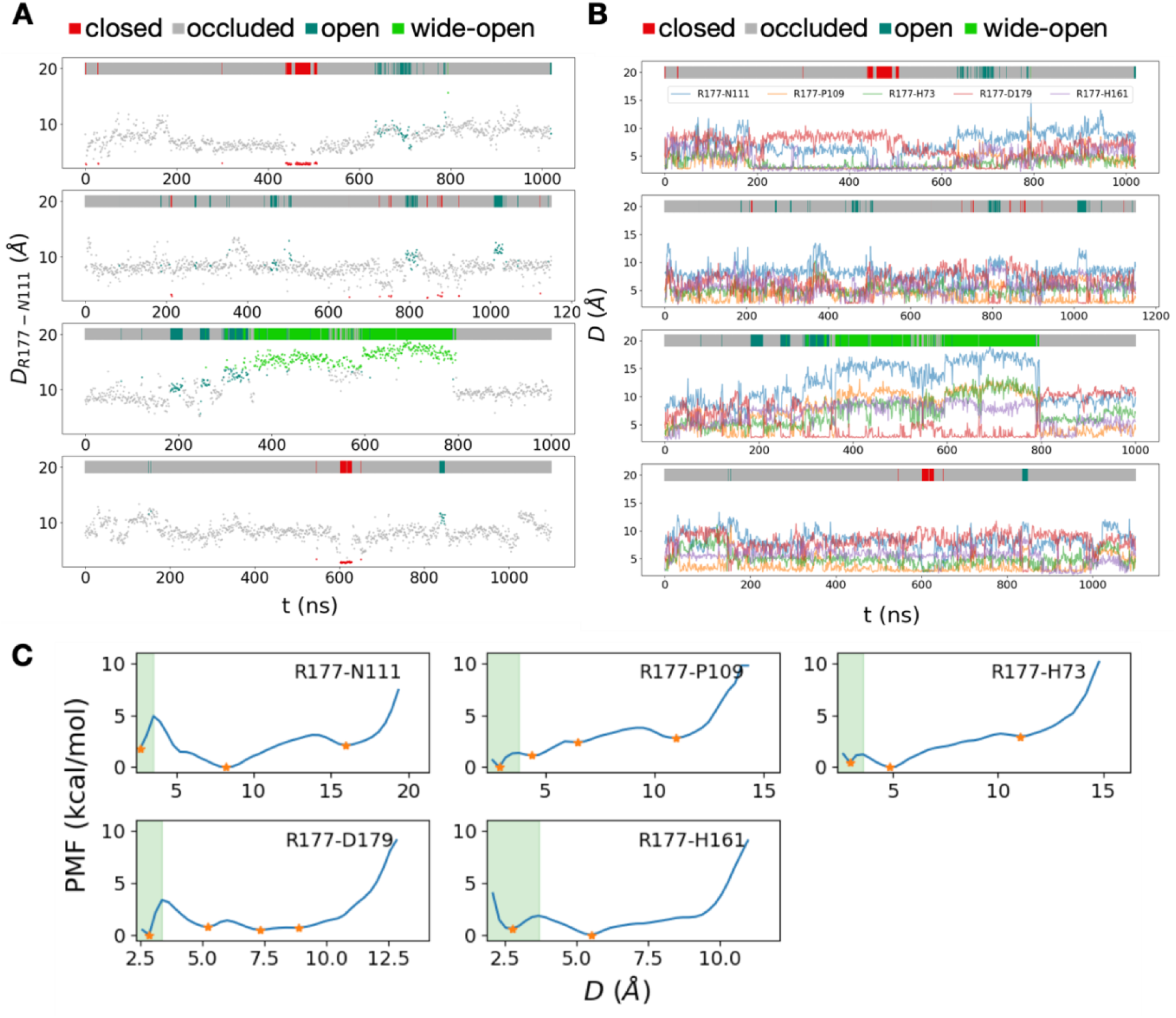
(Panels A-B) Distances between R177 and its hydrogen bonding partners at 1 ns intervals in four unbiased MD trajectories. The colors of individual data points and the bars at the top of each panel indicate the four states, closed (red), occluded (grey), open (acqua) and wide-open (green). (Panel A) R177-N111. (Panel B) Distances between the sidechain of R177 and the sidechains of N111, H161 and D179 and the backbones of HIC73 and P109. (Panel C) Free energies vs. distances between R177 and residues with which it forms hydrogen-bonds. Orange stars indicate energy minimums and green shaded areas depict regions where hydrogen bonds or salt bridges form between corresponding residue pairs. The energy profiles depend on the distinct chemical properties of the residues and their surrounding environments. For example, the R177-N111 distance has three distinct basins in the energy landscape. The minimum around 2.5 Å, is associated with the hydrogen bond between R177 and N111. The other basins at 8.5 Å and 16.0 Å represent interactions that stabilize the system such as hydrogen bonds with other residues such as P109 and D179.

Trajectories of unbiased MD simulations show how the protein fluctuates among these states: closed state Δ multiple occluded states Δ open states Δ wide open states **(Figs. 5A-B)**.

#### Closed state

In the closed state, the distance between R177 and N111 (*D_R177-N111_*) is < 3.53 Å and R177 also forms a hydrogen bond with the backbone of P109. **Fig. 4A** shows these hydrogen bonds and **Fig. 4B** shows that molecular surfaces occlude the channel for phosphate release.

#### Open state

The distances of R177 to its potential hydrogen-bonding partners are *D_R177-P109_* >3.8 Å, *D_R177-HIC73_* >3.57 Å, D_R177-H161_ >3.66 Å and D_R177-D179_ <3.40 Å **(Fig. 4E)**. Without a hydrogen bond between N111 and R177, N111 is reoriented to hydrogen bond with E107 and is located further from R177, accounting for much of the separation. Thus, in the open state R177 forms a salt bridge with D179 but does not form hydrogen bonds with P109, H73 or H161 and leaves open the pore for phosphate to escape **(Fig. 4F and Supplementary Video 2)**.

#### Wide-open state

In this configuration with *D_R177-N111_* >13.7 Å, R177 forms a salt bridge with D179. In both the open and wide-open states the back door is open, creating a pore larger than the size of phosphate between the interior and exterior of the actin subunit **(Fig. 4F)**.

In our simulations, P_i_ consistently moved towards the side chain of R177 as it exited from the protein and was fully released (**Supplementary Video 2)**. This is likely an electrostatic interaction between negatively charged P_i_ and positively charged R177 as shown by the charge distribution in **Fig. 4F**. This transient interaction accounts for the third and smallest cluster of P_i_ in Fig. 3A and 3B.

#### Occluded states

We define occluded states as all configurations of the R177 hydrogen-bonding network other than the three states defined above. These include states with one or more rapidly fluctuating hydrogen-bonds between R177 and HIC73, P109 and/or H161 but not an electrostatic interaction with D179 (**Fig. 4C-D**). N111 forms a hydrogen bond with E107 rather than R177. Analysis of the molecular surfaces shows that the pathway for phosphate between the interior cavity and the exterior of the actin subunit is blocked **(Fig. 4D)**.

During both types of MD simulations starting in the occluded state, the occluded state is the most heavily populated. However, the subunits in available cryo-EM structures of filaments are either in the closed or open states **(Table 2)**.

**Supplementary Video 3** illustrates the three stages previously discussed. The video was assembled from snapshots from a trajectory of one WT-MetaD simulation as explained in supplemental materials. As a consequence of the bias imposed in WT-MetaD simulations, this series of frames differs from the actual-time course of the events. However, the video accurately represents the tendency for the system to occupy predominantly in stage 1, with P_i_ bound to Mg^2+^ (frames 1-208). After P_i_ dissociates from Mg^2+^ (frame 209), the release channel mostly stays in the occluded state. At frame 224 the molecule transitions to the open state before P_i_ escapes through the backdoor gate. The persistence of the occluded state and the fast transition dynamics of the backdoor gate means that the gate has a secondary role in blocking P_i_ release.

### Why are occluded states missing from cryo-EM structures?

The “rapid” cooling during cryo-EM sample preparation is slow enough for room temperature structures to relax to other stable states as observed by Bock and Grubmüller (41) in their simulations of cooling structure ensembles. To investigate whether the R177 hydrogen-bonding network might reconfigure during preparation for cryo-EM, we changed the temperature during unbiased simulations. Starting in the closed state, lowering the temperature from 322 K to 77 K over 40 ns or alternating the temperature between 322 K to 77 K (a common strategy for finding states with minimal energy) did not change the hydrogen bonds of R177 **(Figs. 6B-C)**. However, starting with the occluded state, lowering the temperature reduced the distance between R177 and N111 **(Fig. 6D)**, indicating a transition towards the closed state. These simulations suggest that the closed state is more stable than the occluded state during freezing, which may explain the absence of the occluded state in frozen cryo-EM samples.

**Figure 6.**
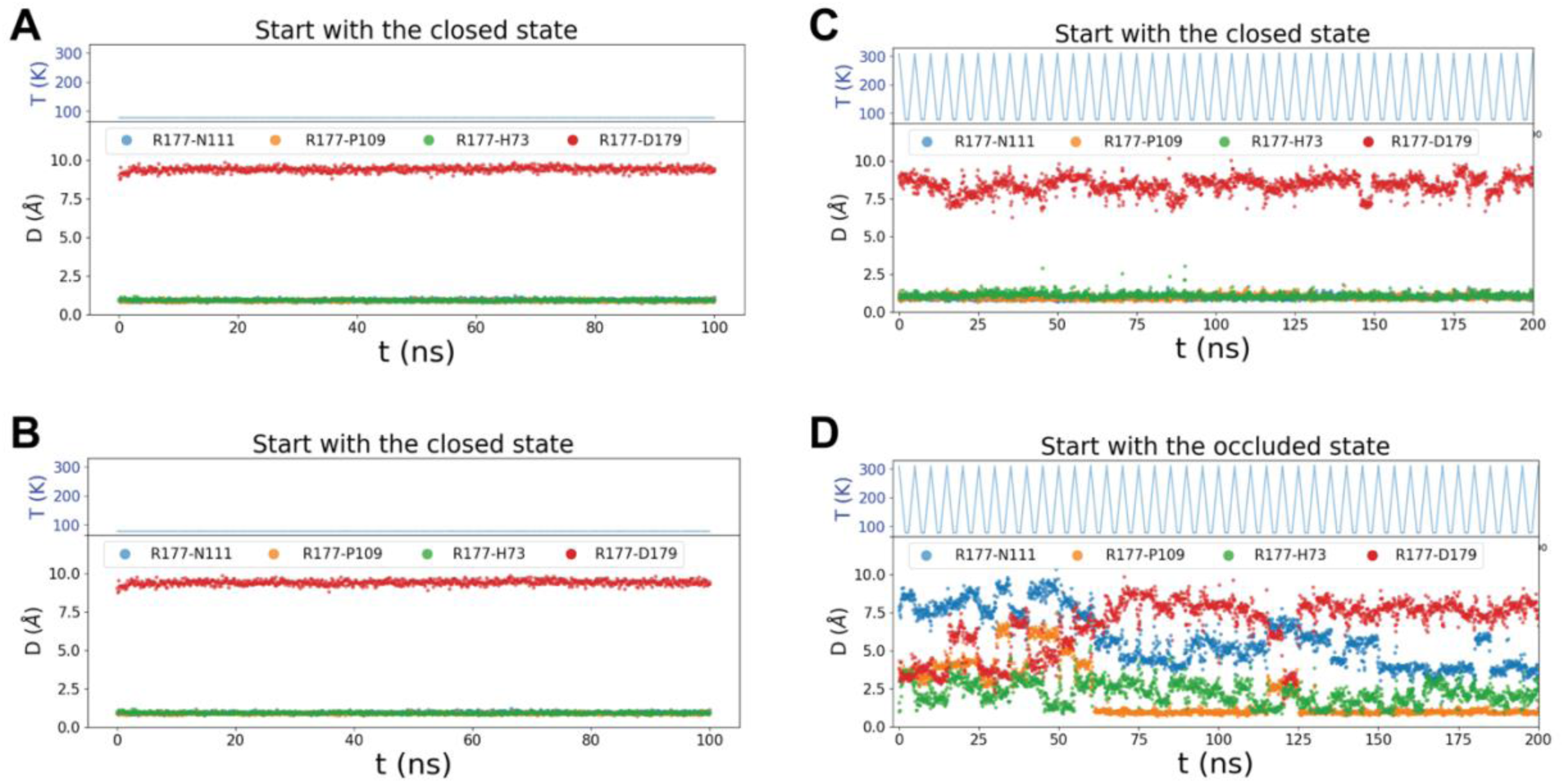
Unbiased MD simulations employing various temperature schemes. (Panel A) Starting with the closed state the temperature was maintained at 77 K throughout the simulation. (Panel B) Starting with the closed state the temperature gradually decreased from 322 K to 77 K over the initial 40 ns and remained constant at 77 K thereafter. (Panels C-D) The temperature was alternated periodically using a linear decrease from 322 K to 77 K over 2 ns, followed by a constant temperature of 77 K for 1 ns, before a linear increase back to 322 K over 2 ns. (Panel C) Starting with the closed state. (Panel D) Starting with the occluded state. The backdoor gate remained closed state with the temperature schemes in panels A-C and tended from the occluded state toward the closed state in panel D.

### Additional occluded and wide-open states

Both types of MD simulations revealed a dynamic hydrogen-bonding network in the phosphate release channel giving rise to multiple metastable states that may be difficult to define based solely on cryo-EM structures and by inspection of the simulation data. To search for overlooked or more obscure states we used a machine learning method called state predictive information bottleneck (SPIB) (42) to identify metastable states automatically from the high-dimensional MD trajectories.

SPIB builds upon the Information Bottleneck (IB) principle (43, 44) and aims to derive a concise yet informative representation of an input source with respect to a target variable. Notably, SPIB incorporates a dynamic learning process during training, enabling the acquisition of a discrete-state representation of the system. We calculated the pairwise distances between atoms from R177 and other residues (listed in **Table 3**) from the unbiased trajectories for use as the input for SPIB to learn the discrete-state representation.

**Table 3.**
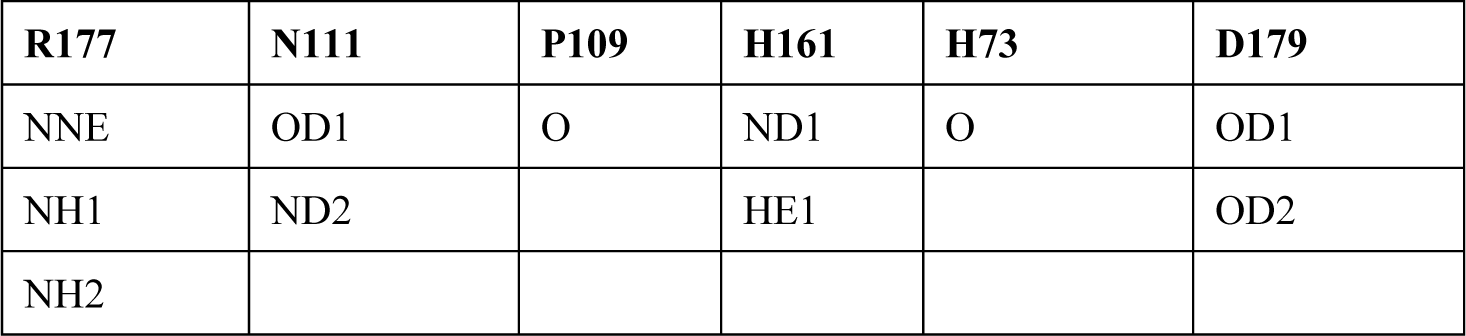
List of atoms that form the network of H-bonds around the phosphate release channel.

**Table 4.** Transition probabilities between states. Each row has the probabilities of moving from the state represented by that row to other states represented by the columns. Self-transition probabilities are light gray and the dominant inter-state transition probabilities and blue.

|  | closed | occluded | open | wide-open |
| --- | --- | --- | --- | --- |
| closed | 0.96 | 0.04 | 0.0 | 0.0 |
| occluded | 0.0010 | 0.9883 | 0.008 | 0.002 |
| open | 0.0 | 0.116 | 0.857 | 0.028 |
| wide-open | 0.0 | 0.021 | 0.017 | 0.961 |

SPIB defined not only the single closed and open states we identified based in cryo-EM structures, but also identified multiple occluded and wide-open states rather than just one of each **(Supplementary Table. S1)**. None of the occluded substates has a hydrogen bond between R177 and N111 or a salt bridge of R177 with D179, consistent with our definition of the occluded state. All occluded substates have a hydrogen bond between R177 and P109. In one occluded substate R177 has additional hydrogen bonds with HIC73 and H161. None of the wide-open substates has hydrogen bonds of R177 with HIC73, P109, N111 or H161, but two of the three substates have a salt bridge between R177 and D179.

### Transitions among the four states of the gate

During long unbiased MD simulations, hydrogen bonds between R177 and formed and dissociated on a nanosecond time scale **(Figs. 5A, 5B)**. Transitions from the closed state to the open state and back always involve the occluded state as the intermediate. Thus, the closed state only transits to the occluded state. The occluded state is the most heavily populated and rarely transits to the closed or open state, so it dominates in blocking phosphate release. The open state is four times more likely to transit to the occluded state than the wide-open state, while the wide-open state has similar probabilities of transiting to the open and the occluded state.

### Simulations starting with a structure with resolved waters

The recent cryo-EM structure of the ADP-P_i_-actin filament at 2.22 Å resolution (Oosterheert, 2022 (28); PDB 8A2S) has clear density for waters in the active site. Four water molecules coordinate with Mg^2+^ at distances around 2.1 Å (Supplementary Fig. S8A). Unbiased and WT-MetaD simulations starting with the PDB 8A2S and 6DJN structures were similar.

The water-Mg^2+^ coordination numbers fluctuated over a range in unbiased MD simulations, with an average coordination number of 3.4 for the structure based on 6DJN and 3.6 for the structure based on 8A2S (Supplementary Fig. S8B). For both systems, the initial coordination numbers were smaller than the average values, suggesting a tendency for the water molecules around P_i_ to relax to a more compact structure during the simulation.

In both systems phosphate formed similar networks of interactions with surrounding residues (Supplementary Fig. S8A). This pattern includes the formation of hydrogen bonds with S14 and Q137, alongside occasional hydrogen bonds or salt bridges with nearby residues such as A108, D154, and H161.

Importantly, WT-MetaD simulations with both starting structures gave very similar P_i_ release energy barriers (Fig. S8C). In particular, the calculated PMF as a function of the separation of P_i_ and Mg^2+^ had a negligible difference in barrier height of less than 0.5 kcal/mol.

## Discussion

The release of phosphate (P_i_) from ADP-P_i_-actin filaments is a critical step in the polymerization cycle of actin. Assembly of purified ATP-actin monomers starts with unfavorable formation of short oligomers, followed by rapid elongation, especially at the fast-growing barbed end. Bound ATP is hydrolyzed rapidly (k_-_ = 0.3 s^-1^) without changing the conformation of the subunit and phosphate dissociates slowly (k_-_ = 0.002 s^-1^) accompanied by subtle conformational changes. This process reaches a steady state where the overall addition and loss of actin monomers is balanced, with subunits preferentially added at one end of the filament (barbed end) and lost at the other (pointed end) owing to phosphate dissociating more rapidly at the pointed end than the barbed end (45). Under some conditions phosphate also dissociates faster from subunits at the barbed end than the interior of the filament (14). During disassembly in the absence of free actin monomers, ADP-actin subunits dissociate more rapidly from barbed ends than pointed ends. Thus, phosphate dissociation is the transformative event in the polymerization cycle, promoting disassembly and binding of disassembly proteins such as cofilin. In cells, regulatory proteins can control these dynamic assembly and disassembly cycles during motility and shape changes.

We used unbiased and enhanced (biased) all-atom MD simulations to investigate the mechanism of P_i_ release from internal ADP-P_i_ subunits in actin filaments at the atomic level. The results, including explicit quantitative PMF profiles, were similar with simulations starting with the computationally solvated 6DJN structure and the higher resolution 8A2S structure having densities for water. We confirmed suggestions from previous studies proposing a "back door" route for P_i_ release (6, 30) and characterized the energy barriers along the pathway. Our WT-MetaD and unbiased MD simulations revealed that P_i_ release is a three-step process involving two energy barriers and four main states of the release pathway.

### Dissociation of phosphate from Mg^2+^

As originally proposed by Wriggers and Schulten (30), the high barrier associated with dissociation of phosphate from Mg^2+^ determines the overall slow rate of the reaction. Initially, the change in coordination between Mg^2+^ and water molecules, along with the conformational changes near the nucleotide-binding site, destabilizes the interaction between Mg^2+^ and P_i_, leading to the dissociation of P_i_. This process makes the largest contribution to the overall PMF for the pathway and only occurs after the disruption of the interaction between phosphate and Q137.

Experimentally, after hydrolysis, inorganic phosphate dissociation from interior subunits proceeds at a rate of 0.002 s^-1^ (16) at room temperature. Assuming that subunit flattening associated with assembly increases ATP-hydrolysis 40,000 fold (7, 8), and subunits at the barbed end are progressively flattened as they acquire neighboring subunits (38), most phosphate release reactions from a growing filament occur in interior subunits at the barbed end. This slow rate implies an energy barrier of ∼21 kcal/mol, assuming a transmission coefficient of unity in the Eyring transition state theory rate constant equation. In our simulations, an energy barrier of 20.5 ± 0.6 kcal/mol separates the P_i_-Mg bound state from the P_i_-Mg unbound state, the highest energy barrier during the release process. The agreement between this quantity and the experimentally derived energy barrier provides strong evidence that the breaking of coordination between Mg^2+^ and P_i_ is the rate-limiting step during P_i_ release. Other interactions in the active site accompany the dissociation of P_i_ from Mg^2+^, including the rearrangement of several residues to disrupt their interactions with P_i_ and the coordination between Mg^2+^ and water molecules. These factors collectively contribute to the high energy barrier in the PMF profile for the P_i_ release.

We were unable to simulate the reverse process, rebinding of P_i_ to the active site, due to the high entropy barrier for a free P_i_ that must be overcome. Therefore, we followed work using similar methods to study ligand unbinding (46-49), and refer to the free energy profiles obtained from these simulations as an approximate potential of mean force, PMF. Although specialized methods can be used to overcome this limitation (50), further research is needed to explore this possibility in-depth.

The rates of P_i_ release from ATPase proteins can differ by orders of magnitude. For example, phosphate dissociates from ADP-P_i_-myosin-VI at 0.04/s (51), about 20 times faster (0.002/s) than core subunits of Mg-ADP-P_i_ actin filaments but similar to P_i_ release from the ends of Mg-ADP-P_i_ actin filaments (45). If we assume the strength of P_i_-Mg^2+^ interaction is the same for different ATPases, interactions of P_i_ with protein residues must contribute to the different release rates. We note that a 20-fold difference in release rates from actin and myosin-VI corresponds to an energy barrier difference less than 1.8 kcal/mol, similar to the energy of one H-bond.

In our simulations of actin filaments and in simulations of myosin-VI (52), phosphate release involves a stepwise hydration of Mg^2+^. In our simulations, the coordination number of water molecules around Mg^2+^ fluctuates on a relatively short time scale, but the release of P_i_ only occurs when four water molecules coordinate with the magnesium ion. Notably, the simulation of myosin-VI revealed that rotating the orientation of P_i_ to weaken its interaction with Mg^2+^could potentially accelerate the release rate by a factor of 1000, shifting from millisecond to microsecond timescales (52). This result also underscores the pivotal role of P_i_-Mg^2+^ interaction in the P_i_ release process.

### Internal movements of phosphate and fluctuations in the gate controlling phosphate release

After dissociation from Mg^2+^, P_i_ diffuses through a channel near HIC73 in subdomain 1 (SD1) of the actin subunit, where it encounters a second, lower energy barrier associated with the transition of the R177 gate from the occluded state to the open state to release phosphate. The exit gate is closed or occluded most of the time, but when the protein fluctuates into the open state, P_i_ moves towards the R177 gate and dissociates from the protein. During four 1 µs unbiased MD simulations, the R177 gate opened multiple times (**Supplementary Fig. S5**).

We did not observe multiple transitions between all metastable states in the MD time courses in **Fig. 5**, suggesting that none of these simulations achieved complete sampling of conformational probability of the backdoor gate. These dynamics are also reflected in the relatively low energy barrier for the final stage of the release process on the PMF profile (**Fig. 1D**). Therefore, opening and closing the backdoor gate may affect the relative rate of P_i_ release from the actin subunits in different confirmations along the filament but does not determine the overall slow rate of P_i_ release.

The occluded states revealed by our MD simulations are most prevalent in filaments at room temperature. In the occluded state, R177 does not hydrogen bond with N111, its partner in the closed state. Instead, R177 participates in a rapidly rearranging network of hydrogen bonds stabilizing conformations that block the pathway for phosphate to dissociate from actin. Interestingly, these states have not been captured in cryo-EM structures. However, our simulations suggest that the occluded state may rearrange to the closed state during freezing.

### Comparisons with other work

Oosterheert et al. (53) used multiple methods to investigate the mechanism of phosphate release from ADP-P_i_-actin filaments. Using an enhanced sampling protocol, they identified pathways of phosphate release with gates formed by R177 or R183. Consistent with Oosterheert et al.’s mutagenesis and biochemical experiments, our WT-MetaD simulations showed that the R177 backdoor pathway is the dominant pathway for P_i_ release. Oosterheert et al. also observed the breaking of the R177 and N111 hydrogen bond in shorter unbiased MD simulations, but not the full opening of R177 and P_i_ release except in steered MD simulations where they drove the configuration of the central subunit to the barbed-end configuration. Thus, the fully open state has a lower probability than other metastable states, but we show that it is accessible by unbiased MD.

Oosterheert et al. also reported a 3.6 Å resolution reconstruction of the barbed end of an ADP-actin filament with the backdoors on the internal subunits closed by hydrogen bonds between R177 and N111. They also noted a repositioning of the H161 side chain in the barbed end as well as the N111S mutant. Given our observation that the forming of a hydrogen bond between H161 and Q137 always accompanies the disruption of P_i_-Mg^2+^ interaction, we propose an alternative hypothesis for the accelerated release of P_i_ in the mutant: the altered position of H161 could enhance its propensity to form a hydrogen bond with Q137, thereby destabilizing the Q137-P_i_ interaction and weakening the overall P_i_ binding affinity.

### Summary

Our study provides new insights into the mechanism of P_i_ release from actin filaments at an atomic level as well as a quantitative identification of the rate limiting step. Our findings may also have important implications for understanding how actin filaments function in various cellular processes such as cell motility and cytokinesis. The results further demonstrate the ongoing value of enhanced sampling methods combined with all-atom MD simulations and machine learning for investigating complex biomolecular processes at the atomic level.

## Methods and Materials

### System and simulation setup

The system was constructed using a 5-mer section of interior subunits from the ADP-Pi 13-mer simulations performed in Zsolnay PNAS 2020 with an updated Charmm36m force field (54) and phosphate parameters generated with cgenff (55). To generate the 13-mer system in Zsolnay PNAS 2020, 13 ADP-P_i_-F-actin subunits (PDB 6DJN) were patterned according to the rise and twist parameters reported in Chou & Pollard (PNAS 2019). The system was solvated using the autosolvate plugin in VMD (56) to add a TIP3P water box with a minimum separation of 11 Å between the protein and the periodic boundary. The autoionize plugin in VMD was used to neutralize the system and bring the solvent to a salt concentration of 100 mM KCl. The system was minimized, heated, and equilibrated with a series of constraints detailed in Zsolnay PNAS 2020. The system underwent a 320 ns unconstrained equilibration.

To construct the 5-mer system, five subunits from the interior of the 13-mer system (subunits 5 to 9) were selected after the unconstrained equilibration of 320 ns. All water molecules within 8 Å of the subunits were included in the selection. Autosolvate and autoionize were used to fill the remainder of the solvation box to guarantee a separation of 11 Å between the protein and the periodic boundary and a neutral system with 100 mM KCl. The initial structure was subjected to energy minimization and the minimization was terminated when the maximum force is smaller than 1000.0 kJ/mol/nm. After energy minimization, the system was equilibrated with an NPT ensemble for 10 ns with a 2 fs integration step. The Parrinello-Rahman barostat (57) and Bussi-Parrinello velocity rescaling thermostat (58) were used to simulate the constant NPT ensemble at temperature 310 K and pressure 1 bar. The conformations of the terminal subunits at both ends changed during this process.

To prepare the simulation with the PDB 8A2S structure, we first used MODELLER (59) to fill in missing residues. Hydrogens are added by the CHARMM-GUI input generator tool (60). We added hydrogens to phosphate oxygens using the strategy of Mugnai and Thirumalai in their simulation of P_i_ release from myosin-VI, avoiding the oxygen atom of P_i_ closest to Mg^2+^ (52). Similar to the preparation of the initial structure from the 13-mer, we kept the water molecules at the site from the PDB structure and used VMD Autosolvate and Autoionize to fill the remainder of the solvation box. To further relax the system, especially refine the orientations of P_i_ and water molecules near Mg^2+^, we performed 100 ns constant NVT simulations with constraints added on heavy atoms of protein, ADP, Mg^2+,^ and phosphorus of P_i_, enabling the P_i_ and water molecules surrounding Mg^2+^ to adopt stable orientations or positions. Since only the phosphorus atom of the phosphate was restrained, the oxygen and hydrogens of the phosphate were free to move into a stable configuration. Regardless of which oxygen atoms received the two hydrogens, the relaxed structures always resembled the one shown in Supplementary Fig. S9A, with one hydrogen atom located at the oxygen closest to ADP and the other hydrogen located at the oxygen near Q137.

All simulations used to study the release process were performed with the same constant NPT ensemble. Simulations were performed by using GROMACS 2020.4 compiled with PLUMED 2.4. The data generated from the simulations were analyzed by VMD (56), PyMol (61-63) and Python library MDanalysis (64, 65), and an in-house Python script.

We define the four subdomains of an actin molecule as: subdomain 1 (SD1): residues 1 to 32, 70 to 144, and 338 to 375; (SD2): residues 33 to 69; (SD3): residues 145 to 180, 270 to 337; (SD4): residue 181 to 269. The center of mass of the Cα atoms of each subdomain were used to compute a dihedral angle SD2-SD1-SD3-SD4.

We consider the existence of hydrogen bonds between a donor and acceptor pair when the distance between the electronegative atoms is smaller than 3.6 Å. Due to the observation of fluctuating hydrogen bond angles during simulations, the angle is not utilized as a strict criterion for the presence of hydrogen bonding.

### Metadynamics simulations

The timescale for the release of P_i_, which occurs over several minutes, is orders of magnitude longer than the timescale accessible through MD simulations, which typically run on the microsecond scale at best. To overcome this limitation and sample this rare event, we used WT-MetaD (34-37) to accelerate the release process. Unlike a conventional metadynamics (MetaD) simulation, which has convergence issues, WT-MetaD decreases the height of Gaussian bias according to the existing biasing potential and has demonstrated convergence properties (35).

The choice of reaction coordinates (i.e., collective variables, CVs) are critical for the success of MetaD simulations. MetaD can enhance the sampling of rare events only when chosen CVs provide a good representation of the process of interest. In this study, we employed two CVs to perform our WT-MetaD simulation: the distance between Mg^2+^ and the center of mass (COM) of P_i_, denoted as d _Mg-Pi,_ and the distance between the COM of the backbone R177 and the COM of P_i._ denoted as d_R177-Pi_. The selection of these CVs was informed by both general intuition and previous studies of phosphate release. The cryo-EM structure indicates that the position of Mg^2+^ does not change after the release of P_i_. Therefore, the distance between Mg^2+^ and P_i_ can be considered a good CV as it indicates how far P_i_ is from its binding position. The equilibrium structure from an unbiased MD simulation presents a cavity inside the central actin subunit (Fig. 3). The cavity has the shape of a channel with the side chain R177 situated at its end in the closed state. It is natural to expect that after breaking the coordination with Mg^2+^, P_i_ can diffuse in the cavity. Previous studies based on structure also indicate the important role of R177 in the release process. These factors motivated us to use the P_i_-R177 distance, which is defined as the COM of P_i_ and the COM the backbone of R177, as the second RC. To make the RC less noisy, we only considered the backbone of R177 and avoided accounting for the fluctuation of its side chain.

To estimate the free energy profile (i.e., PMF), we utilized the approach suggested in Ref. (36) where the samples with microscopic coordinate *R* from WT-MetaD are each associated with a reweighting factor:

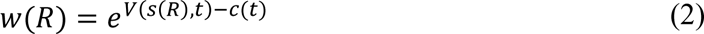

where *V*(*x*) is the bias *c*(*t*) is usually referred to as a time-dependent bias offset. The term *c*(*t*) can be estimated “on the fly” in the MetaD using the form:

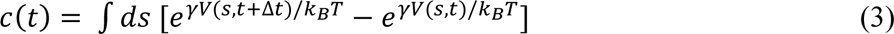

where Δ*t* is the time interval between successive bias depositions. The unbiased estimation of an observable can be calculated as:

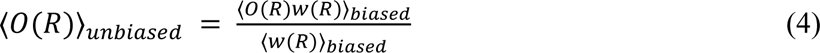

Next, the PMF can be calculated by *F*(*x*) = −*k*_*B*_*T ln*〈*δ*(*x* − *x*(*R*))〉_*unbiased*_. Note that since here *x* can be any observable and is not necessary to be the reaction coordinate *s*, the free energy profiles of other variables of interest can also be obtained.

The choice of MetaD parameters of a WT-MetaD simulation includes the biasing factor, biasing frequency and bias height and the variance of the Gaussian bias. In our simulations, the biasing factor was set to be 10, the bias height was set to be 1 kcal/mol and the biasing frequency is set to be 5000 MD steps. These parameters were chosen to ensure that the biasing potential was not added too aggressively while still being able to accelerate the release process. The variances of Gaussian bias were set to be 0.05 Å and 0.5 Å separately for d_Pi-Mg_ and d_Pi-R177_ and these values were determined by half of the variance of these variables in a 50 ns unbiased MD simulation.

## Acknowledgements

Research reported in this publication was supported by the National Institute of General Medical Sciences (NIGMS) of the National Institutes of Health (NIH) under award numbers R01GM063796 to GAV and R01GM026132 to TDP. The content is solely the responsibility of the authors and does not necessarily represent the official views of the National Institutes of Health. YW gratefully acknowledges the support of Chicago Center for Theoretical Chemistry Fellowship and the Eric and Wendy Schmidt AI in Science Postdoctoral Fellowship. The authors thank the University of Chicago Research Computing Center and the NIH-funded Beagle-3 computer (NIH award 1S10OD028655-01) for computational resources.

