## Supplemental Material for "Mechanism of Phosphate Release from Actin Filaments"

**Supplemental Materials**

**Supporting Information Text**

**Water coordination number**

The water coordination number of $Mg^{2+}$ is calculated using a switching function and has the form:

$$c = \frac{1}{3}\sum_{i\in W} \frac{1-\left(\frac{r_i}{r_0}\right)^6}{1-\left(\frac{r_i}{r_0}\right)^{12}} \qquad \text{(S1)}$$

Here $r_0$ is set to be 3 Å and the summation is over all atoms of water molecules.

**Distance between residues**

H-bonds can form between distinct atom pairs from two residues during the course of a simulation. To characterize the H-bonding network between residues, we monitored the smallest distance between atoms that participate in forming hydrogen and electrostatic bonds. The distance between two residues I and J reported in our study is defined by:

$$D_{I-J} = \min_{i\in I, j\in J} d_{i-j} \qquad \text{(S2)}$$

Where $d_{i-j}$ is the distance between atom i and j, which belong to residue *I* and *J* separately and participate in forming H-bonds (listed in Table 2).

**Transition probability between different conformations of the backdoor gate**

To estimate the transition probability between states, we first labeled each frame according to our definition of the open, wide-open states, and occluded states. Then, we went through four unbiased trajectories and counted the number of transitions between states. Next, the number of transitions was recorded as an element of a transition matrix *T*. To obtain probabilities, we normalized the matrix by dividing each element in the transition matrix by the total count of transitions from the corresponding state. This step ensures that the matrix represents a valid probability distribution.

**State Predictive Information Bottleneck**

SPIB utilizes a neural network with an encoder and decoder structure to effectively cluster the high-dimensional MD trajectories into metastable states. The loss function used to train the network in the SPIB method is based on the Information Bottleneck (IB) principle. The IB principle postulates that the desired representation should use minimal information from the input to predict the target. Therefore, the neural network is trained to extract only the essential information from past trajectories, which is crucial for accurately predicting the future state of the system. This objective can be phrased as the task of optimizing a probabilistic encoder p($\mathbf{z}|\mathbf{x}$) and a probabilistic decoder p($\mathbf{y}|\mathbf{z}$) to maximize the following objective function:

$$\boldsymbol{L_{IB}} = \boldsymbol{I}(\boldsymbol{z},\boldsymbol{y}) - \boldsymbol{\beta I}(\boldsymbol{X},\boldsymbol{z}) \tag{S3}$$

Where $\boldsymbol{I}(\cdot)$ is the mutual information between two random variables. This objective function is designed to force the encoder to compress the information from input feature $\boldsymbol{X}$ and the decoder then uses the information constrained in the latent representation $\boldsymbol{z}$ to predict some preassigned feature $\boldsymbol{y}$. which, in SPIB, is the future state label learned during the training.

The direct optimization of the information bottleneck is impractical as the calculation of mutual information is computationally expensive. Therefore, following the approach in Ref. 43 in the main text, SPIB employs a variational lower bound on the original objective function. This lower bound, used as the loss function for training, is given by the formula:

$$\begin{aligned} L_{IB} \geq & \frac{1}{N}\sum_{n=1}^{N}\int dz[-p(z \mid X^n)\log q(y^n \mid z)] \\ & -\beta\frac{1}{N}\sum_{n=1}^{N}\int dz\left[p(z \mid X^n)\log\frac{p(z \mid X^n)}{r(z)}\right] + H(y) \end{aligned} \tag{S4}$$

Where $q(\boldsymbol{y}|\boldsymbol{z})$ and $r(\boldsymbol{z})$ are variational approximations to the true probability distributions $p(\boldsymbol{y}|\boldsymbol{z})$ and $p(\boldsymbol{z})$, respectively.

SPIB adopts an iterative approach to learn the number and state labels dynamically. It is based on the assumption that if a configuration is in state *i* at a certain time then, after a time delay, it is most likely to remain in state *i*. Accordingly, in SPIB, the state label of a configuration $\boldsymbol{X}$ is updated to match the state label predicted by the decoder of the trained network.

In summary, the SPIB is first initialized with a randomly initialized state label. Then, the learning process occurs with an iteration between network training and state label update according to the prediction of the trained network. The learning process is converged when all the state labels do not change in the update. During the training process, the network learns a discrete-state representation, focusing specifically on the motion associated with state-to-state transitions.

For our specific implementation, we selected a deep neural network as the encoder to effectively capture the nonlinear relationship among the input variables. The training dataset comprised 24 pairwise distances between atoms involved in forming the H-bond network around R177. Default hyperparameters were employed, and a time delay of 3 ns was chosen to filter out fast dynamic.

**Initial Structure Preparation for constrained simulations**

We used Steered-MD (SMD) to create the initial structures for constrained simulations, with all subunit dihedral angles being driven to -3°. The harmonic restraints $\boldsymbol{V_r}$ applied in SMD have the form:

$$V_r = k\big(\theta - \theta_0(t)\big)^2, \tag{S5}$$

where $\boldsymbol{k}$ is the force constant and $\boldsymbol{\theta_0}$ is the time-dependent center of the harmonic SMD restraint. We first applied harmonic restraints with a force constant of 5000 kJ/mol/radian$^2$ on the dihedral angles of all subunits. The center of harmonic restraints moves to -3.01 radians in 10 ns with a constant velocity. Then we fixed the centers of harmonic restraints and linearly decreased or increased the force constant between 5000 kJ/mol/radian$^2$ and 100 kJ/mol/radian$^2$ with a 20 ns period. This relaxation process helped the system better find the local minimum. Figure S6 shows how the center and the force constant change during the simulation of preparing the initial structure. After the SMD simulations, the restraint on the central subunit was removed, and a 100 ns simulation was initialized to further equilibrate the system. The initial structure was chosen from the frame with the central unit dihedral angle being closest to -3° during the last equilibrium phase.

**Interactions between $P_i$ and two ATPases: actin filaments and myosin-VI**

ATPase enzymes release $P_i$ with a wide range of rates. For instance, release from myosin-VI releases $P_i$ at 0.04/s, 400 times faster than $P_i$ release from Mg-ADP-$P_i$ actin filaments. A high energy barrier in our simulations suggests that the dissociation of $P_i$ from $Mg^{2+}$ is the rate limiting step. To investigate if the interaction network between $P_i$ and the surrounding residues might explain the difference of energy barriers in ATPase enzymes, we compared the interaction network in actin and myosin-VI (Supplementary Fig. S8) in the crystal structure of Mg-ADP-$P_i$ bound actin filament (PDB code 6DJN) and ADP-$P_i$-$Mg^{2+}$ bound myosin-VI (PDB code 4PFP). We highlight all the protein-$P_i$ atom pairs with distances fewer than 4 Å. We observed 6 such pairs in the Mg-ADP-$P_i$ bound actin filament and 5 in the ADP-$P_i$-$Mg^{2+}$ bound myosin-VI, implying a potentially stronger interaction and a more stabilized bound state of $P_i$ in the actin filament. However, the strength of these interactions is influenced by multiple factors, including the chemical properties and distances of the atom pairs. Additionally, the fluctuation of the structure also affects the stability of the interaction. Therefore, a simulation study of phosphate dissociation from ADP-$P_i$-$Mg^{2+}$ bound myosin would be informative but falls outside the scope of our current focus on actin filaments.

### Supplemental Tables

**Table S1.** SPIB analysis of unbiased MD simulations: learned states and distances (Å) between R177 and interacting residues. The listed distances represent the average values calculated across the ensemble of each SPIB state. Red font indicates distances short enough to form hydrogen or electrostatic bonds. See Fig. S1 for time courses.

| SPIB state | R177-N111 SC | R177-P109 C=O | R177-H161 SC | R177-H73 C=O | R177-D179 SC | Gate |
|---|---|---|---|---|---|---|
| Closed | 2.8 | 2.8 | 3.9 | 3.1 | 8.70 | Closed |
| Open | 10.3 | 5.64 | 6.5 | 6.14 | 6.0 | Open |
| Occluded 1 | 8.2 | 4.2 | 5.5 | 4.9 | 7.0 | Occluded |
| Occluded 2 | 6.0 | 2.8 | 2.9 | 3.0 | 7.5 | Occluded |
| Occluded 3 | 9.5 | 3.9 | 4.7 | 6.4 | 10.5 | Occluded |
| Occluded 4 | 8.2 | 3.4 | 5.6 | 7.8 | 9.5 | Occluded |
| Wide-open 1 | 13.6 | 9.0 | 8.2 | 7.6 | 4.5 | Wide-open |
| Wide-open 2 | 14.7 | 9.9 | 8.5 | 7.6 | 3.0 | Wide-open |
| Wide-open 3 | 16.8 | 11.3 | 8.5 | 11.2 | 3.1 | Wide-open |

**Supplemental Figures**

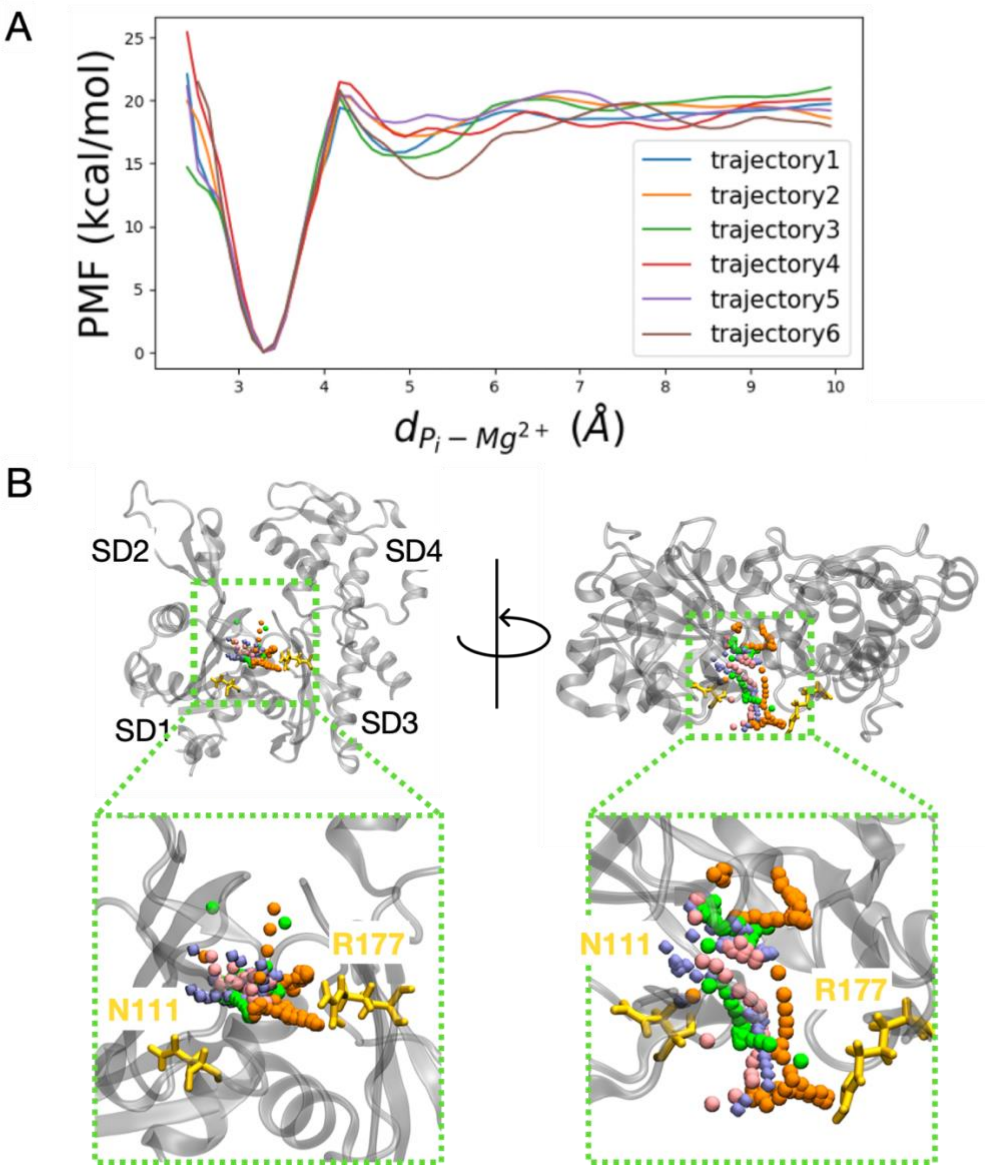


**Fig. S1.** Pathways of $P_i$ release in independent WT-MetaD simulations. (Panel A) The potential of mean force (PMF) profiles from six independent WT-MetaD trajectories. The low variance across the simulations suggests the release pathway is robust. (Panel B) Four trajectories of phosphate release from actin subunit A2 are made by plotting the positions of the phosphorus in each frame. For the purpose of visualization, the frames are not spaced uniformly in time. R177 and N111 (yellow) indicate the exit of the "backdoor" pathway.

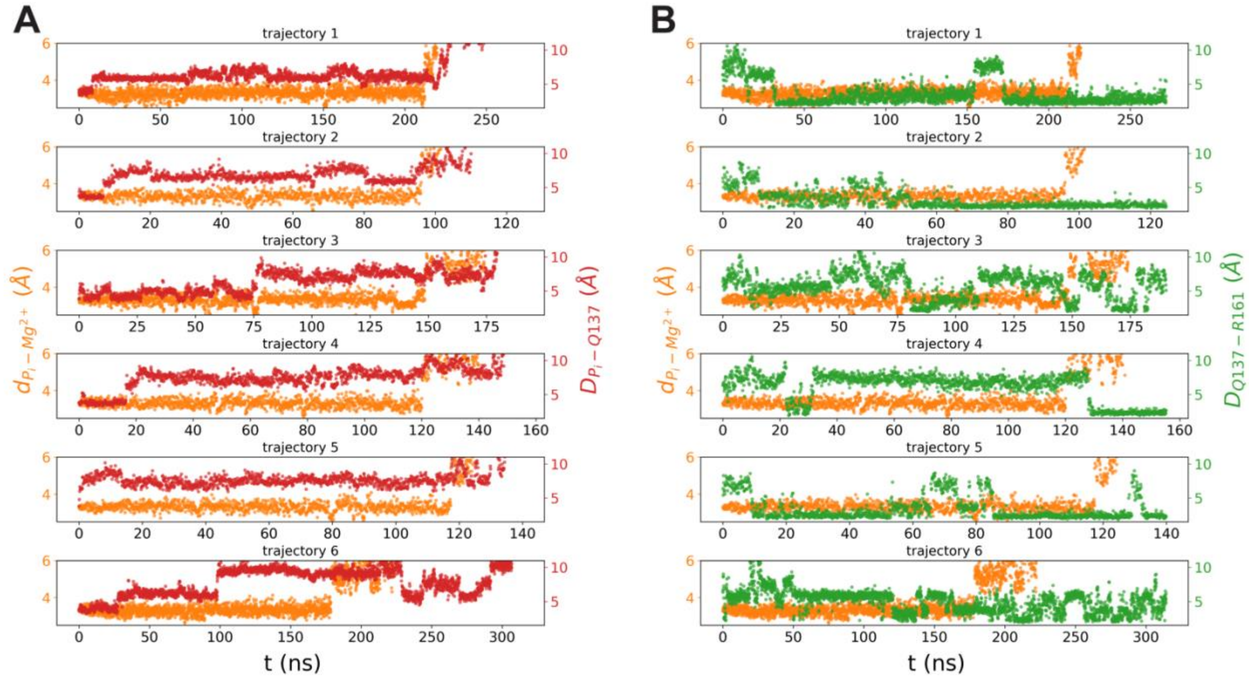


**Fig. S2.** Time courses from six independent WT-MetaD simulations plotting distances between: (Panel A) phosphate and $Mg^{2+}$ (orange), phosphate and Q137 (red); (Panel B) phosphate and $Mg^{2+}$ (orange), and between the atoms of Q137 and H161 that can form a hydrogen bond (green). Phosphate separates from $Mg^{2+}$ only after Q137 reorients away from $P_i$. Following its reorientation away from $Mg^{2+}$, Q137 can form a hydrogen bond with the side chain of H161. See Fig. 2D, E for visualization of the rearrangement of Q137.

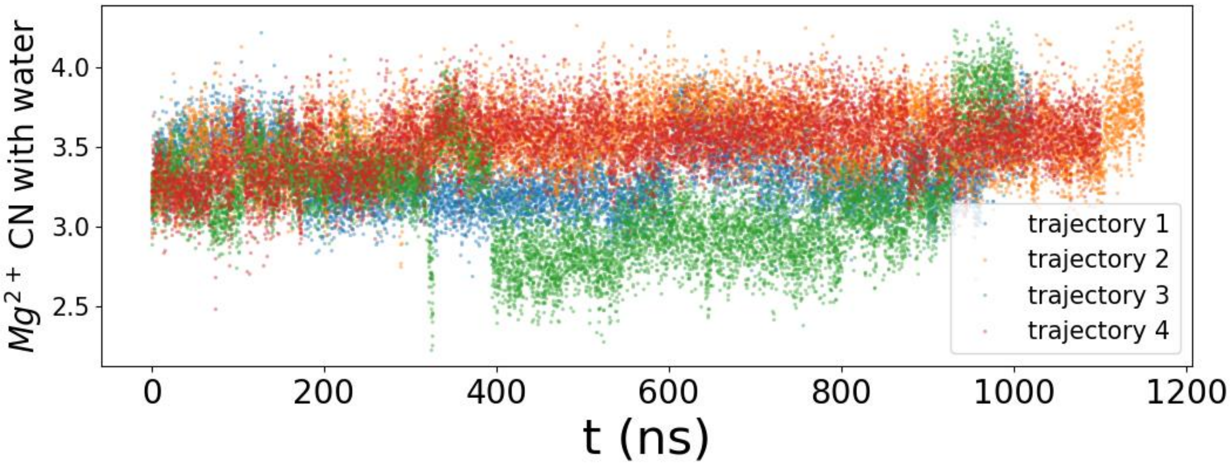


**Fig. S3.** Time courses from three independent unbiased MD simulations of the number of waters coordinated with $Mg^{2+}$. The number fluctuates between 2.3 to 4 over time. See Fig. 2B,C of the main text for visualization of the waters in the active site.

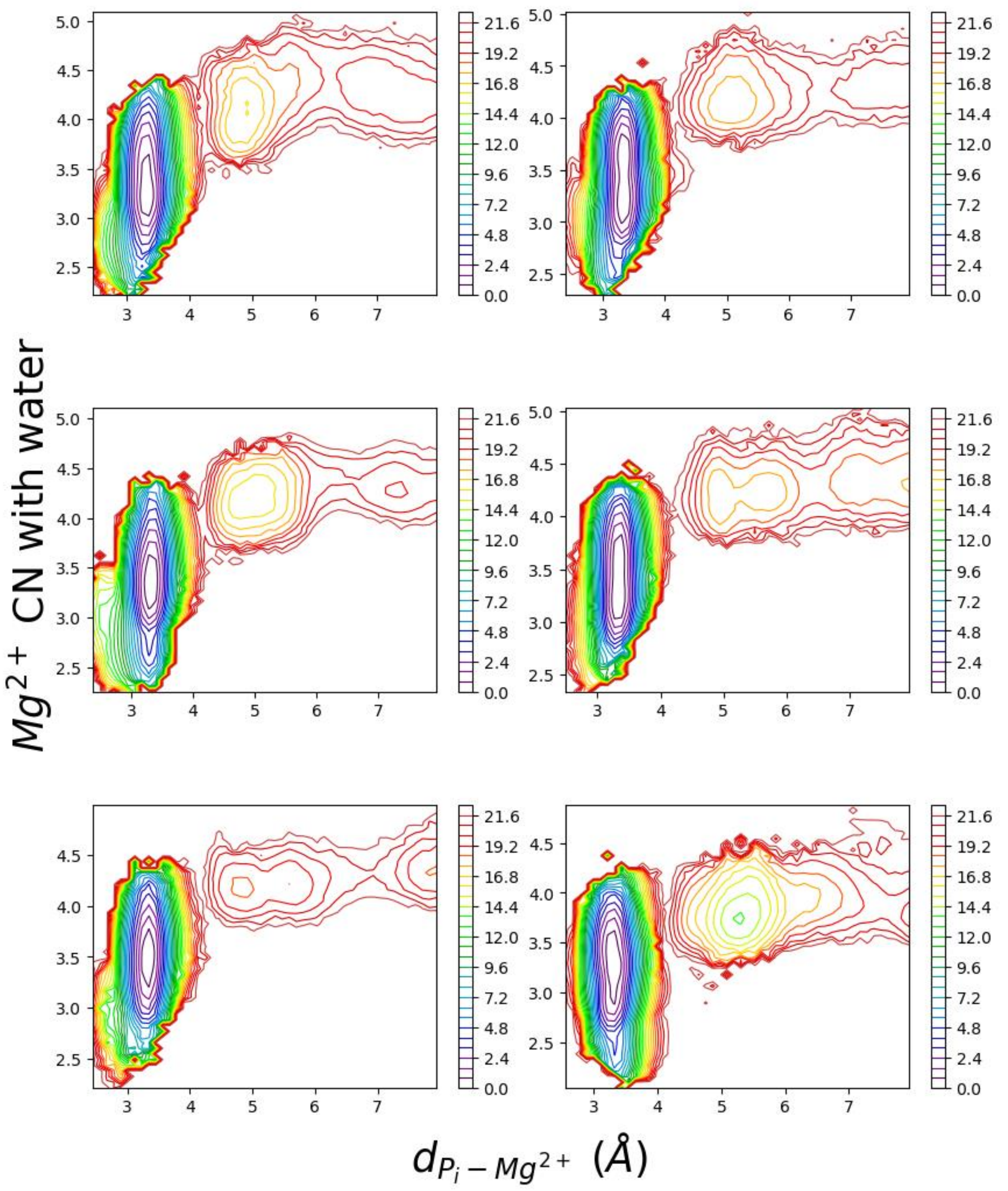


**Fig. S4.** Two-dimensional plots of potential of mean force (PMF; i.e., colored coded free energy contours in kcal/mole) from six independent WT-MetaD trajectories as a function of the number of waters coordinated with $Mg^{2+}$ and the distance between $P_i$ and $Mg^{2+}$. Phosphate separates from $Mg^{2+}$ only after four waters coordinate with $Mg^{2+}$. The upper left panel is the same as Fig. 2A.

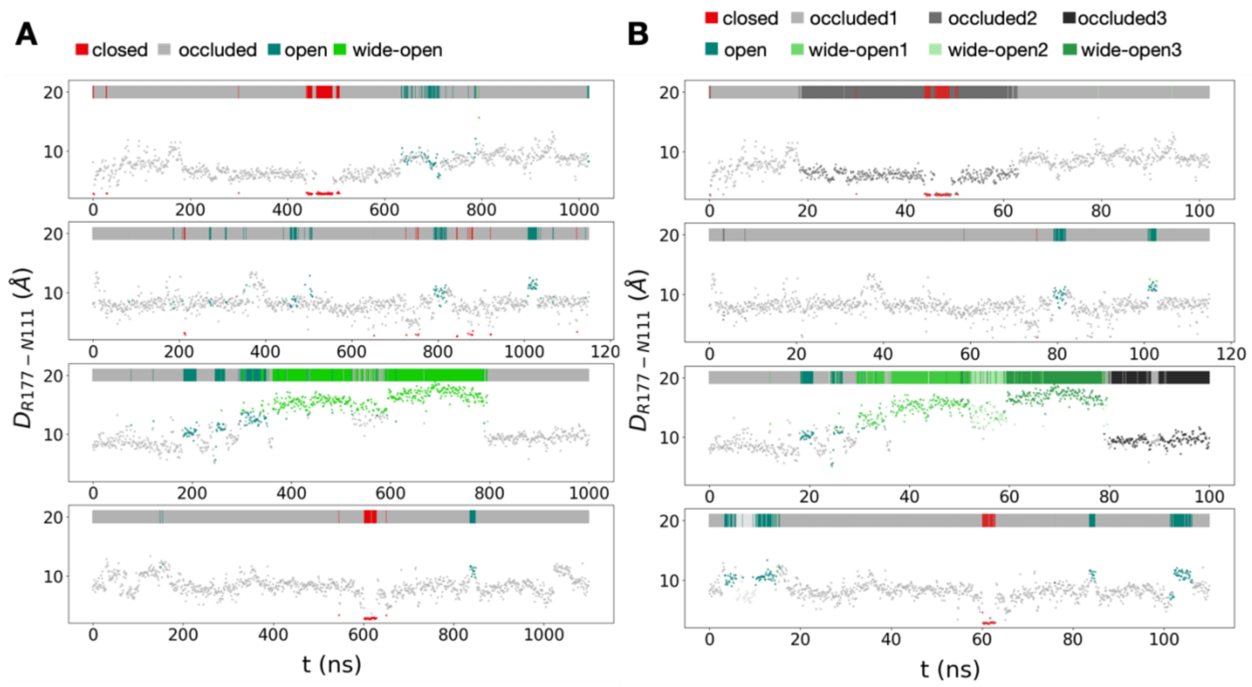


**Fig. S5.** Distances between R177 and N111 in four unbiased MD trajectories with states color-coded according to (Panel A) our definitions and (Panel B) a machine learning method called state predictive information bottleneck (SPIB) to identify metastable states automatically from the high-dimensional MD trajectories. SPIB identified substates of the occluded and wide-open states. See Table S1 for the interactions of R177 in the substates.

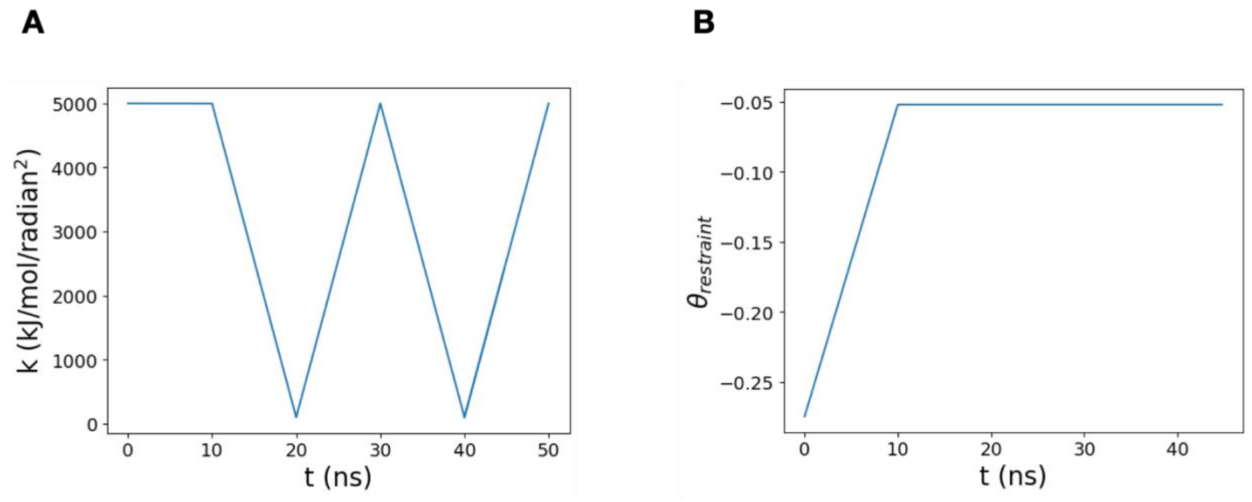


**Fig. S6.** (Panel A) Variation of the force constant $k$ and (Panel B) the displacement of the center $\theta_{restraint}$ of the harmonic potential applied on the subunits to drive them to a flattened state. In the first 10 ns, the center of harmonic potential gradually approached the target dihedral angle (-3°) with a constant velocity and the force constant was kept at a constant of 5000 kJ/mol/radian$^2$. Subsequently, the center of the harmonic potential was fixed, and its force constant oscillated linearly between 5000 kJ/mol/radian$^2$ and 100 kJ/mol/radian$^2$ a period of 20 ns.

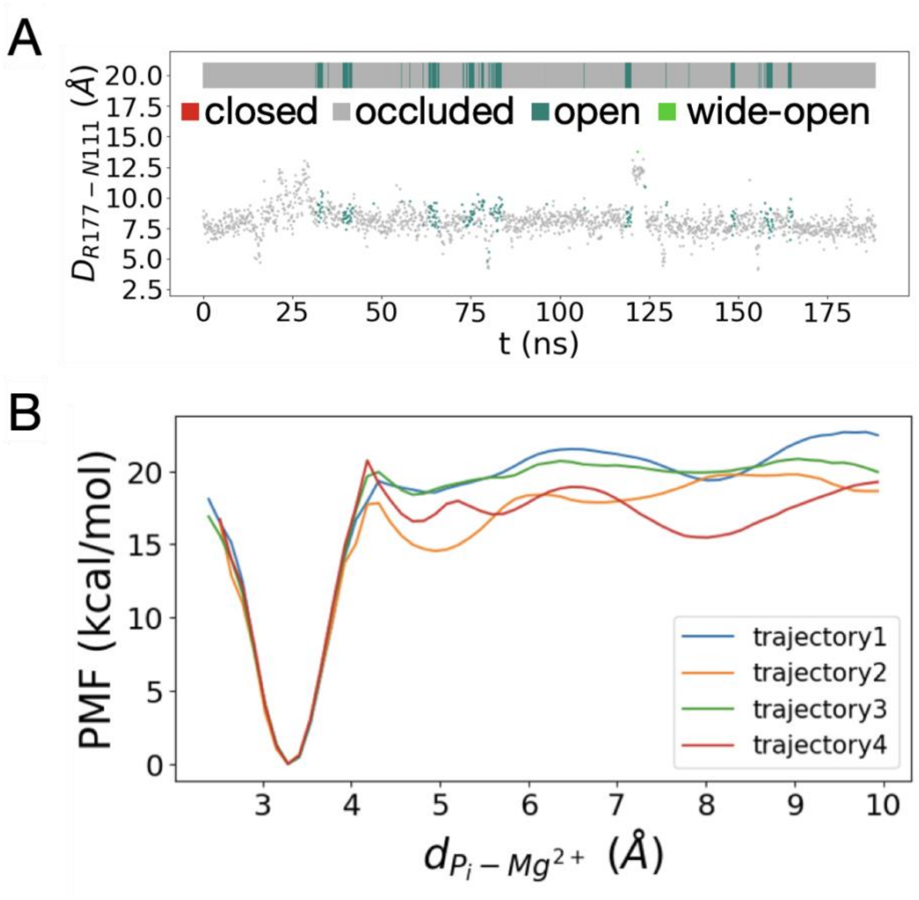


**Fig. S7.** Simulations employing harmonic restraints to preserve the flattened state of the terminal subunits. (Panel A) Distances between R177 and its hydrogen bonding partners at 0.1 ns intervals in an unbiased MD simulation, with data points colored by their backdoor conformations. (Panel B) The potential of mean force (PMF) profiles from four independent WT-MetaD simulations with restrained actin subunits.

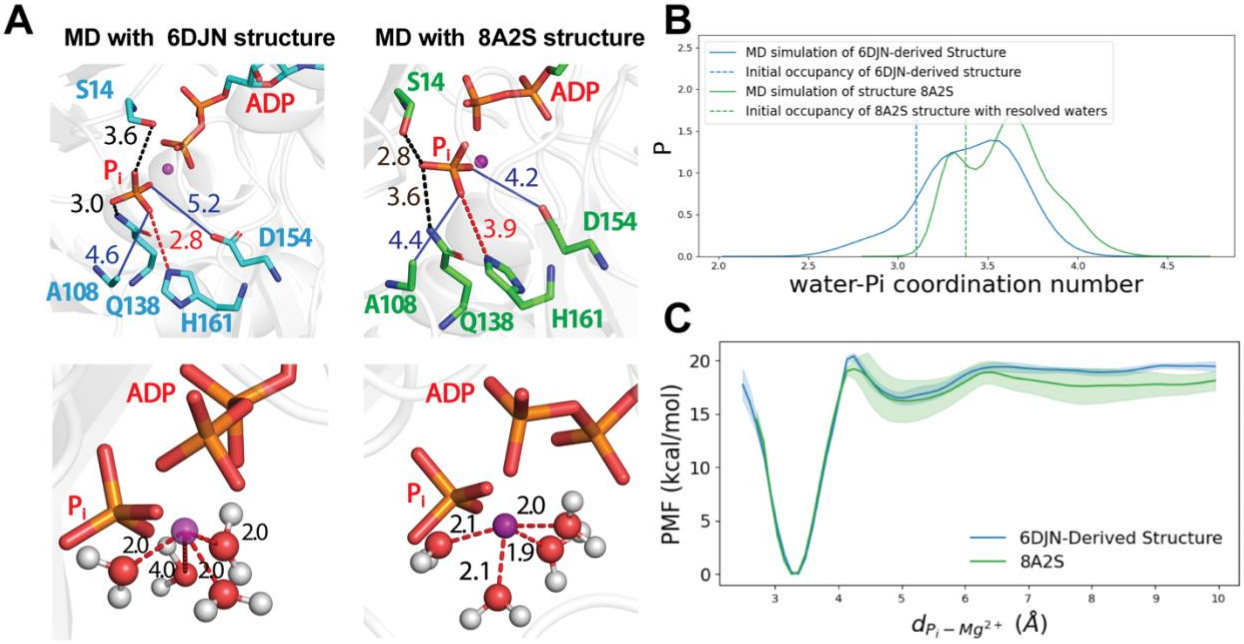


**Fig. S8.** Comparison of MD simulations with structure derived from PDB Structures 6DJN (cryo-EM structure of Mg-ADP-$P_i$ actin filament at 3.1 Å resolution hydrated by MfD) and 8A2S (cryo-EM structure of Mg-ADP-$P_i$ actin filament with waters at 2.22 Å resolution). (Panel A) Molecular interactions that stabilize the bound state of $P_i$. The left column shows the initial structure derived from 6DJN. The right column shows snapshots of the simulation with structure derived from 8A2S. The upper figures illustrate the interactions between $P_i$ and nearby residues. The lower figures focus on water molecules coordinating with $Mg^{2+}$ ions, with dashed lines indicating the electrostatic interactions between water oxygen atoms and $Mg^{2+}$ ions. (Panel B) Distribution of water-$P_i$ coordination number in unbiased MD, colored by blue for the 6DJN-derived simulation and green for the 8A2S-derived simulation. Dashed lines indicate the water-$P_i$ coordination numbers for the initial structures. (Panel C) PMFs from WT-MetaD simulations with two structures, with standard errors indicated by the shaded regions.

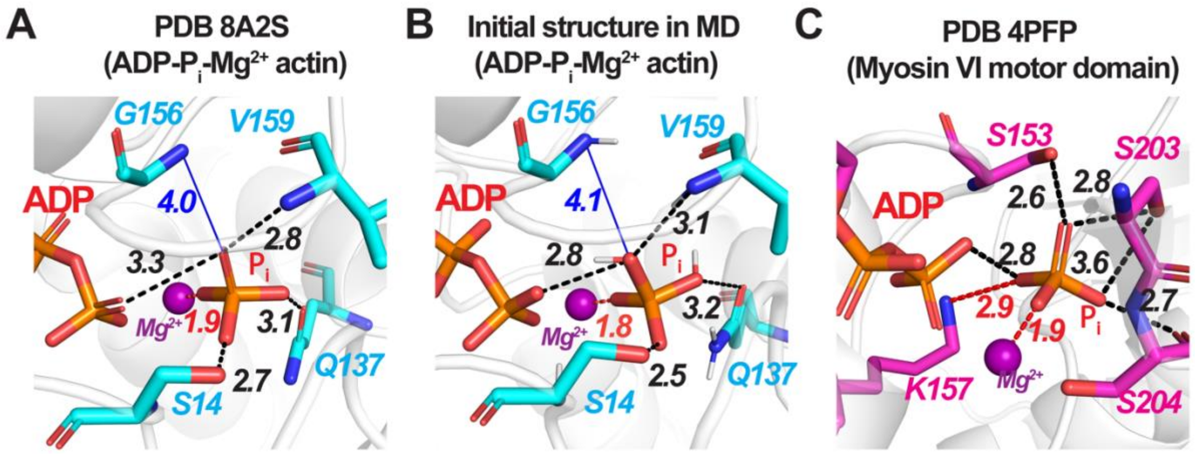


**Fig. S9.** Stick figures comparing interactions between phosphate oxygens and the proteins in ADP-P$_i$-actin and ADP-P$_i$-myosin-VI. Hydrogen bonds are depicted as black dashed line and electrostatic interactions by red dashed lines, while the distances between P$_i$ and other key residues are marked by blue lines. (Panel A) PDB 8A2S structure. (Panel B) Initial structure of ADP-P$_i$-actin in MD simulations (derived from Oosterheert cryo-EM structure PDB 8A2S). (Panel C) The crystal structure of ADP-P$_i$-myosin-VI at 2.32 Å resolution (PDB 4PFP). All the panels focus on the residues in close interaction with P$_i$.

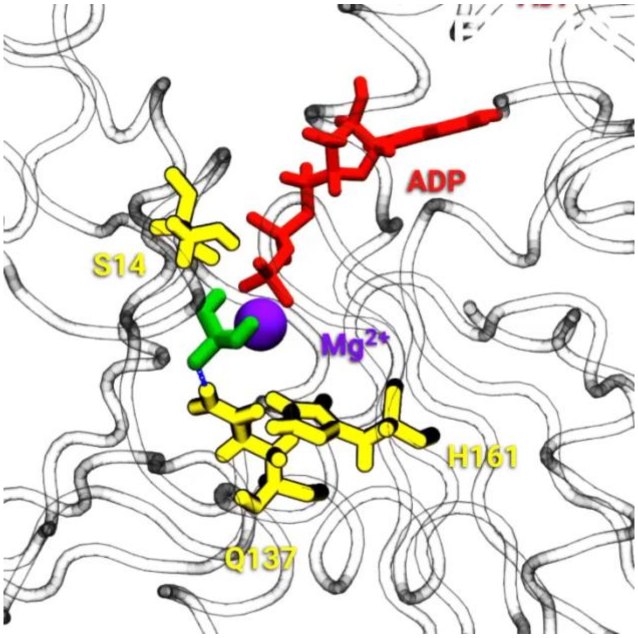


**Supplementary Video 1.** Time lapse video of 45 ns of time from a WT-MetaD MD simulation showing the active site of an actin filament subunit. Licorice representations show the protein backbone and stick figures show ADP (red), phosphate (green) and backbones and sidechains for residues that interact with phosphate (yellow). $Mg^{2+}$ is shown as a purple sphere. Phosphate dissociates from $Mg^{2+}$ only following disruption of its interaction with Q137 and reorientation of Q137 to hydrogen bond with H136.

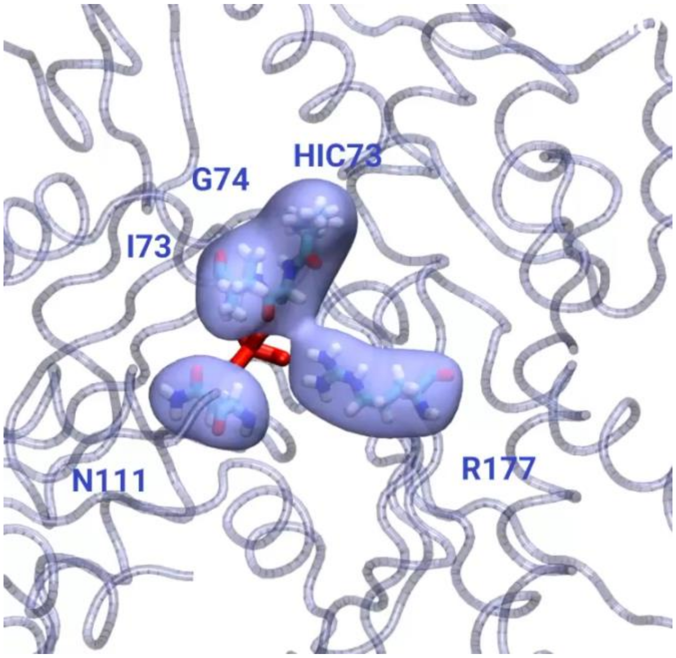


**Supplementary Video 2.** Time lapse video of 5 ns of time from a WT-MetaD simulation showing phosphate release from actin. Residues forming the gate are shown as stick figures inside blue surface representations. Phosphate is a red stick figure.

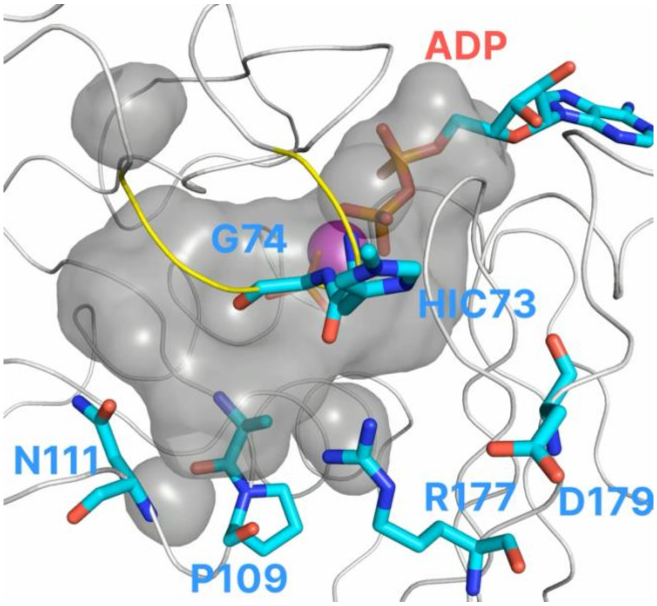


**Supplementary Video 3.** Time lapse video assembled from a WT-MetaD MD simulation that shows the movement of $P_i$ within the cavity and escape through the gate. The frames were recorded every 0.5 ns, and to reduce visual noise, followed by smoothing with a window size of 5. As a consequence of the bias imposed in WT-MetaD simulations, this series of frames differ from the actual-time course of the events. The cavity is shown as a grey surface, $Mg^{2+}$ is shown as a purple sphere, $P_i$, ADP, and residues are shown in the stick representation. The S-loop (residues 70 to 79) is highlighted in yellow. In the first 208 frames, the $P_i$ coordinates tightly with the $Mg^{2+}$ and only moves within its vicinity. After dissociating from $Mg^{2+}$, $P_i$ diffuses locally in the occluded release channel. At frame 224 the molecule transitions to the open state before $P_i$ escapes through the backdoor gate. The fluctuation of the cavity's shape reflects the dynamic nature of the protein’s structure and associated interaction changes.
